# Allosteric conformational ensembles have unlimited capacity for integrating information

**DOI:** 10.1101/2020.12.10.420117

**Authors:** John W. Biddle, Rosa Martinez-Corral, Felix Wong, Jeremy Gunawardena

## Abstract

Integration of binding information by macromolecular entities is fundamental to cellular functionality. Recent work has shown that such integration cannot be explained by pairwise cooperativities, in which binding is modulated by binding at another site. Higher-order cooperativities (HOCs), in which binding is collectively modulated by multiple other binding events, appears to be necessary but an appropriate mechanism has been lacking. We show here that HOCs arise through allostery, in which effective cooperativity emerges indirectly from an ensemble of dynamically-interchanging conformations. Conformational ensembles play important roles in many cellular processes but their integrative capabilities remain poorly understood. We show that sufficiently complex ensembles can implement any form of information integration achievable without energy expenditure, including all HOCs. Our results provide a rigorous biophysical foundation for analysing the integration of binding information through allostery. We discuss the implications for eukaryotic gene regulation, where complex conformational dynamics accompanies widespread information integration.

## INTRODUCTION

Cells receive information in many ways, including mechanical force, electric fields and molecular binding, of which the last is the most diverse and widespread. Elaborate molecular networks have evolved to integrate this information and make appropriate decisions. For molecular binding, the most immediate form of information integration is pairwise cooperativity, in which binding at one site modulates binding at another site. This can arise most obviously through direct interaction, where one binding event creates a molecular surface, which either stabilises or destabilises the other binding event. This situation is illustrated in Fig.1A, which shows the binding of ligand to sites on a target molecule. (In considering the target of binding, we use “molecule” for simplicity to denote any molecular entity, from a single polypeptide to a macromolecular aggregate such as an oligomer or complex with multiple components.) We use the notation *K_i,S_* for the association constant—on-rate divided by off-rate, with dimensions of (concentration)^−1^—where *i* denotes the binding site and *S* denotes the set of sites which are already bound. This notation was introduced in previous work (Estrada et al., 2016) and is explained further in the Materials and methods. It allows binding to be analysed while keeping track of the context in which binding occurs, which is essential for making sense of how binding information is integrated.

**Figure 1:**
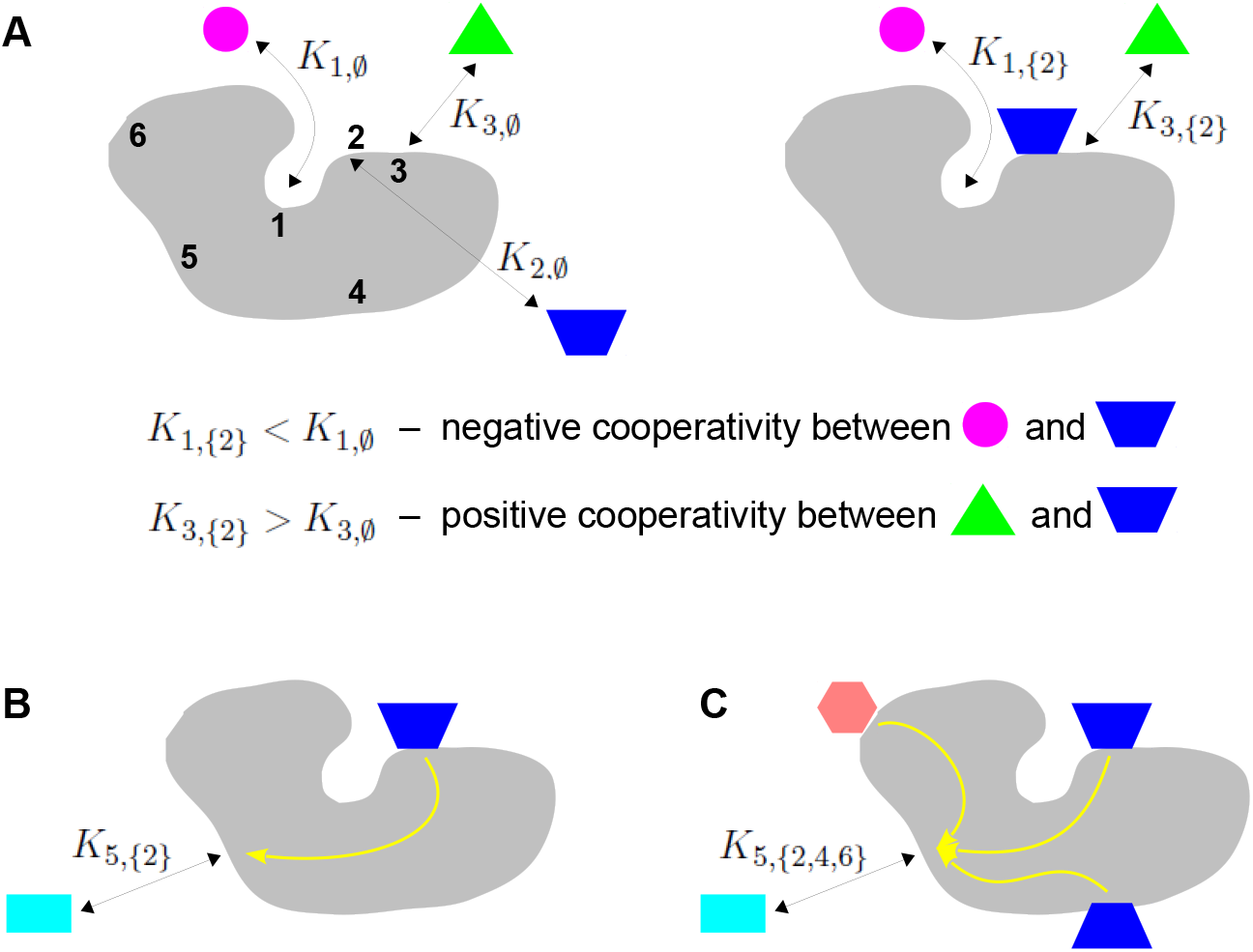
Binding cooperativity. **A.** Pairwise cooperativity by direct interaction on a target molecule (gray). As discussed in the text, the target could be any molecular entity. Left, target molecule with no ligands bound; numbers 1, ···, 6 denote the binding sites. Right, target molecule after binding of blue ligand to site 2. **B.** Indirect long-distance pairwise cooperativity, which can arise “effectively” through allostery. **C.** Higher-order cooperativity, in which multiple bound sites, 2, 4 and 6, affect binding at site 5.

Oxygen binding to haemoglobin is a classical example of integration of binding information, for which Linus Pauling gave the first biophysical definition of cooperativity (Pauling, 1935). At a time when the mechanistic details of haemoglobin were largely unknown, Pauling assumed that cooperativity arose from direct interactions between the four haem groups. He defined the pairwise cooperativity for binding to site *i*, given that site *j* is already bound, as the fold change in the association constant compared to when site *j* is not bound. In other words, the pairwise cooperativity is given by 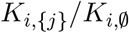, where 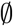 denotes the empty set. (Pauling considered non-pairwise effects but deemed them unnecessary to account for the available data.) It is conventional to say that the cooperativity is “positive” if this ratio is greater than 1 and “negative” if this ratio is less than 1; the sites are said to be “independent” if the cooperativity is exactly 1, in which case binding to site *j* has no influence on binding to site *i*. This terminology reflects the underlying free energy, which is given by the logarithm of the association constant (Materials and methods, Eq.11). Fig.1A depicts the situation in which there is negative cooperativity for binding to site 1 and positive cooperativity for binding to site 3, given that site 2 is bound.

Studies of feedback inhibition in metabolic pathways revealed that information to modulate binding could also be conveyed over long distances on a target molecule, beyond the reach of direct interactions (Changeux, 1961, Gerhart, 2014) (Fig.1B). Monod and Jacob coined the term “allostery” for this form of indirect cooperativity (Monod and Jacob, 1961). Monod, Wyman and Changeux (MWC) and, independently, Koshland, Némethy and Filmer (KNF) put forward equilibrium thermodynamic models, which showed how effective cooperativity could arise from the interplay between ligand binding and conformational change (Koshland et al., 1966, Monod et al., 1965). In the two-conformation MWC model (Fig.2B), there is no “intrinsic” cooperativity—the binding sites are independent in each conformation—and “effective” cooperativity arises as an emergent property of the dynamically-interchanging ensemble of conformations.

**Figure 2:**
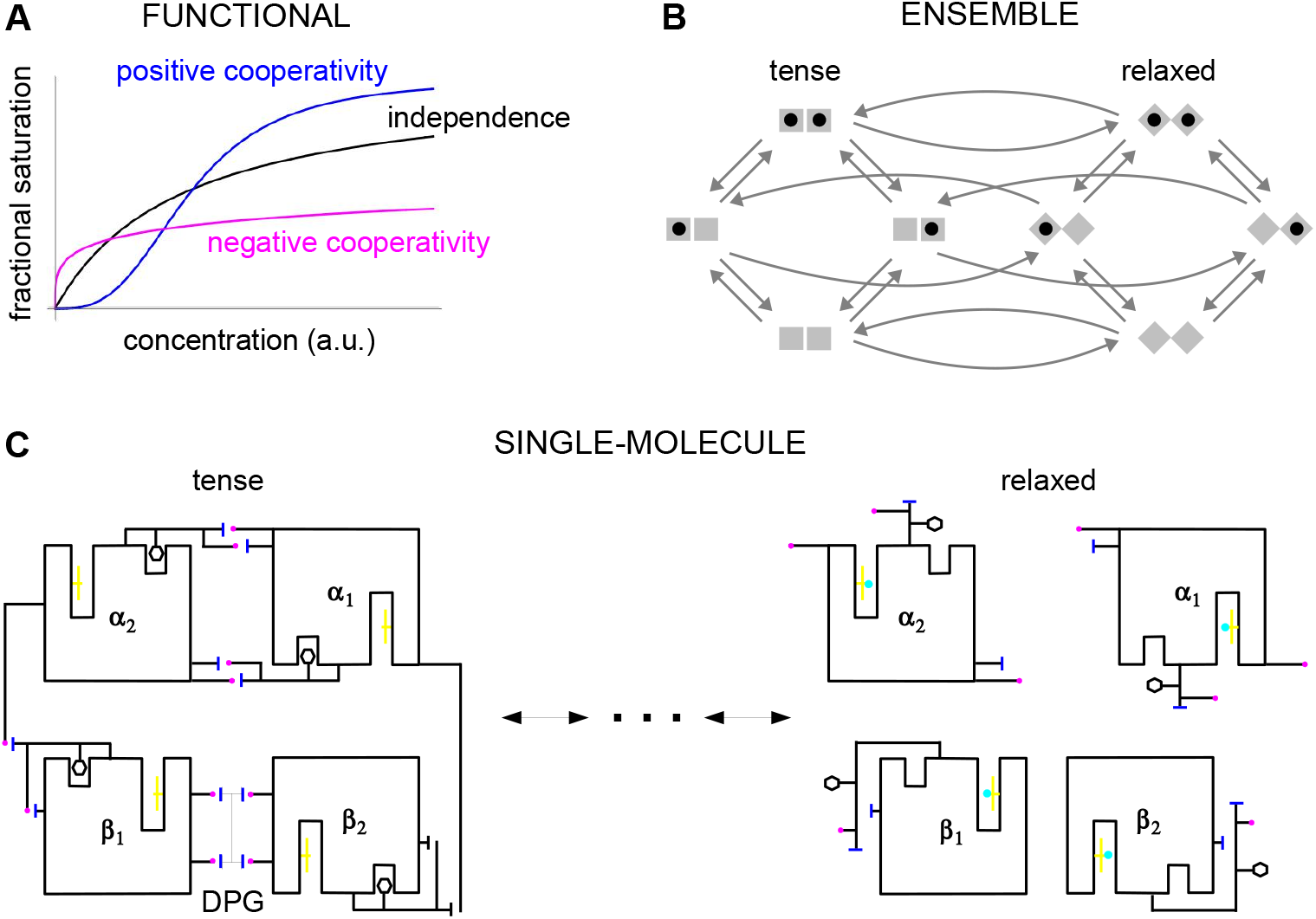
Cooperativity and allostery from three perspectives. **A.** Plots of the binding function, whose shape reflects the interactions between binding sites, as described in the text. **B.** The MWC model with a population of dimers in two quaternary conformations, with each monomer having one binding site and ligand binding shown by a solid black disc. The two monomers are considered to be distinguishable, leading to four microstates. Directed arrows show transitions between microstates. This picture anticipates the graph-theoretic representation used in this paper, as shown in Fig.3. **C.** Schematic of the end points of the allosteric pathway between the tense, fully deoxygenated and the relaxed, fully oxygenated conformations of a single haemoglobin tetramer, *α*_1_*α*_2_*β*_1_*β*_2_, showing the tertiary and quaternary changes, based on (Perutz, 1970, Fig.4). Haem group (yellow); oxygen (cyan disc); salt bridge (positive, magenta disc; negative, blue bar); DPG is 2-3-diphosphoglycerate.

In these studies, the effective cooperativity between sites was not quantitatively determined. Instead, the presence of cooperativity was inferred from the shape of the binding function, which is the average fraction of bound sites, or fractional saturation, as a function of ligand concentration (Fig.2A). The famous MWC formula is an expression for this binding function (Monod et al., 1965). If the sites are effectively independent, the binding function has a hyperbolic shape, similar to that of a Michaelis-Menten curve. A sigmoidal curve, which flattens first and then rises more steeply, indicates positive cooperativity, while a curve which rises steeply first and then flattens indicates negative cooperativity. Surprisingly, despite decades of study, the effective cooperativity of allostery is still largely assessed in this way, through the shape of the binding function rather than a quantitative measure. One of the contributions of this paper is to show theoretically how effective cooperativity can be quantified, thereby placing allosteric information integration on a similar biophysical foundation to that provided by Pauling for direct interactions between two sites.

The MWC and KNF models are phenomenological: effective cooperativity arises as an emergent property of a conformational ensemble. This leaves open the question of how information is propagated between distant binding sites across a single molecule. This question was particularly relevant to haemoglobin, for which it had become clear that the haem groups were sufficiently far apart that direct interactions were implausible. Perutz’s X-ray crystallography studies of haemoglobin revealed a pathway of structural transitions during cooperative oxygen binding which linked one conformation to another (Fig.2C), thereby relating the single-molecule viewpoint to the ensemble viewpoint (Perutz, 1970). These pioneering studies provided important justification for key aspects of the MWC model, which has endured as one of the most successful mathematical models in biology (Changeux, 2013, Marzen et al., 2013).

Allostery was initially thought to be limited to certain symmetric protein oligomers like haemoglobin and to involve only a few, usually two, conformations. But Cooper and Dryden’s theoretical demonstration that information could be conveyed by fluctuations around a dominant conformation (Cooper and Dryden, 1984) anticipated the emergence of a more dynamical perspective (Henzler-Wildman and Kern, 2007). At the single-molecule level, it has been found that binding information can be conveyed over long distances by complex atomic networks, of which Perutz’s linear pathway (Fig.2C) is only a simple example (Schueler-Furman and Wodak, 2016, Kornev and Taylor, 2015, Knoverek et al., 2019, Wodak et al., 2019). These atomic networks may in turn underpin complex ensembles of conformations in many kinds of target molecules and allosteric regulation is now seen to be common to most cellular processes (Nussinov et al., 2013, Changeux and Christopoulos, 2016, Motlagh et al., 2014, Lorimer et al., 2018, Wodak et al., 2019, Ganser et al., 2019). The unexpected finding of widespread intrinsic disorder in proteins has been particularly influential in prompting a reassessment of the classical structure-function relationship, with conformations which may only be fleetingly present providing plasticity of binding to many partners (Wrabl et al., 2011, Wright and Dyson, 2015, Berlow et al., 2018).

However, while ensembles have grown greatly in complexity from MWC’s two conformations and new theoretical frameworks for studying them have been introduced (Wodak et al., 2019), the quantitative analysis of information integration has barely changed beyond pairwise cooperativity. Despite previous hints (Dodd et al., 2004, Peeters et al., 2013, Martini, 2017, Gruber and Horovitz, 2018), little attention has been paid to higher-order cooperativities (HOCs) in which multiple binding events collectively modulate another binding site (Fig.1C). Such higher-order effects can be quantified by association constants, *K_i,S_*, where the set *S* has more than one bound site. The size of *S*, denoted by #(*S*), is the order of cooperativity, so that pairwise cooperativity may be considered as HOC of order 1. For the example in Fig.1C, the ratio, 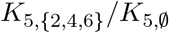, defines the non-dimensional HOC of order 3 for binding to site 5, given that sites 2, 4 and 6 are already bound. The notation used here is essential to express such higher-order concepts.

HOCs were introduced in (Estrada et al., 2016), where it was shown that experimental data on the sharpness of gene expression could not be accounted for purely in terms of pairwise cooperativities (Park et al., 2019a). In this context, the target molecule is the chromatin structure containing the relevant transcription factor (TF) binding sites and the analogue of the binding function is the steady-state probability of RNA polymerase being recruited, considered as a function of TF concentration (Estrada et al., 2016, Park et al., 2019a). The Hunchback gene considered in (Estrada et al., 2016, Park et al., 2019a), which is thought to have 6 binding sites for the transcription factor Bicoid, requires HOCs up to order 5 to account for the data, under the assumption that the regulatory machinery is operating without energy expenditure at thermodynamic equilibrium. An important problem emerging from this previous work, and one of the starting points for the present paper, is to identify molecular mechanisms which are capable of implementing such HOCs.

In the present paper, we show that allosteric conformational ensembles can implement any pattern of effective HOCs. Accordingly, they can implement any form of information integration that is achievable at thermodynamic equilibrium. We work at the ensemble level (Fig.2B), using a graph-based representation of Markov processes developed previously (below). We introduce a general method of “coarse graining”, which is likely to be broadly useful for other studies. This allows us to define the effective HOCs arising from any allosteric ensemble, no matter how complex. The effective HOCs provide a quantitative language in which the integrative capabilities of an ensemble can be specified. It is straightforward to determine the binding function from the effective HOCs and we derive a generalised MWC formula for an arbitrary ensemble, which recovers the functional perspective. Our results subsume and greatly generalise previous findings and clarify issues which have remained confusing since the concept of allostery was introduced. Our graph-based approach further enables general theorems to be rigorously proved for any ensemble (below), in contrast to calculation of specific models which has been the norm up to now.

Our analysis raises questions about how effective HOCs are implemented at the level of single-molecules, similar to those answered by Perutz for haemoglobin and the MWC model (Fig.2C). Such problems are important but they require different methods for their solution (Wodak et al., 2019) and lie outside the scope of the present paper. Our analysis is also restricted to ensembles which are at thermodynamic equilibrium without expenditure of energy, as is generally assumed in studies of allostery. Energy expenditure may be present in maintaining a conformational ensemble, most usually through post-translational modification, but the significance of this has not been widely appreciated in the literature. Thermodynamic equilibrium sets fundamental physical limits on information processing in the form of “Hopfield barriers” (Estrada et al., 2016, Biddle et al., 2019, Wong and Gunawardena, 2020). Energy expenditure can bypass these barriers and substantially enhance equilibrium capabilities. However, the study of non-equilibrium systems is more challenging and we must defer analysis of this interesting problem to subsequent work (Discussion).

The integration of binding information through cooperativities leads to the integration of biological function. Haemoglobin offers a vivid example of how allostery implements this relationship. This one target molecule integrates two distinct functions, of taking up oxygen in the lungs and delivering oxygen to the tissues, by having two distinct conformations, each adapted to one of the functions, and dynamically interchanging between them. In the lungs, with a higher oxygen partial pressure, binding cooperativity causes the relaxed conformation to be dominant in the molecular population, which thereby takes up oxygen; in the tissues, with a lower oxygen pressure, binding cooperativity causes the tense conformation to be dominant in the population, which thereby gives up oxygen. Evolution may have used this integrative strategy more widely than just to transport oxygen and we review in the Discussion some of the evidence for an analogy between functional integration by haemoglobin and by gene regulation.

## RESULTS

### Construction of the allostery graph

Our approach uses the linear framework for timescale separation (Gunawardena, 2012), details of which are provided in the Supplementary Information (Materials and methods) along with further references. We briefly outline the approach here.

In the linear framework a suitable biochemical system is described by a finite directed graph with labelled edges. In our context, graph vertices represent microstates of the target molecule, graph edges represent transitions between microstates, for which the edge labels are the instantaneous transition rates. A linear framework graph specifies a finite-state, continuous-time Markov process and any reasonable such Markov process can be described by such a graph. We will be concerned with the probabilities of microstates at steady state. These probabilities can be interpreted in two ways, which reflect the ensemble and single-molecule viewpoints of Fig.2. From the ensemble perspective, the probability is the proportion of target molecules which are in the specified microstate, once the molecular population has reached steady state, considered in the limit of an infinite population. From the single-molecule perspective, the probability is the proportion of time spent in the specified microstate, in the limit of infinite time. The equivalence of these definitions comes from the ergodic theorem for Markov processes (Stroock, 2014). These different interpretations may be helpful when dealing with different biological contexts: a population of haemoglobin molecules may be considered from the ensemble viewpoint, while an individual gene may be considered from the single-molecule viewpoint. As far as the determination of probabilities is concerned, the two viewpoints are equivalent.

The graph representation may also be seen as a discrete approximation of a continuous energy landscape (Frauenfelder et al., 1991), in which the target molecule is moving deterministically on a high-dimensional landscape in respone to a potential, while being buffeted stochastically through interactions with the surrounding thermal bath. In mathematics, this approximation goes back to the work of Wentzell and Freidlin on large deviation theory for stochastic differential equations in the low noise limit (Ventsel’ and Freidlin, 1970, Freidlin and Wentzell, 2012). It has been exploited more recently to sample energy landscapes in chemical physics (Wales, 2006) and in the form of Markov state models arising from molecular dynamics simulations (Noé and Fischer, 2008, Sengupta and Strodel, 2018). In this approximation, the microstates correspond to the minima of the potential up to some energy cut-off, the edges correspond to appropriate limiting barrier crossings and the labels correspond to transition rates over the barrier.

The linear framework graph, or the accompanying Markov process, describes the time-dependent behaviour of the system. Our concern in the present paper is with systems which have reached a steady state of thermodynamic equilibrium, so that detailed balance, or microscopic reversibility, is satisfied. The assumption of thermodynamic equilibrium has been standard since allostery was introduced (Koshland et al., 1966, Monod et al., 1965) but has significant implications, as pointed out in the Introduction, and we will return to this issue in the Discussion. At thermodynamic equilibrium, we can dispense with dynamical information and work with what we call “equilibrium graphs”. These are also directed graphs with labelled edges but the edge labels no longer contain dynamical information in the form of rates but rather ratios of forward to backward rates. These ratios are the only parameters needed at equilibrium. From now on, in the main text, when we say “graph”, we will mean “equilibrium graph”.

We explain such graphs using our main example. Fig.3 shows the graph, *A*, for an allosteric ensemble, with multiple conformations *c*_1_, ···, *c_N_* and multiple sites, 1, ···, *n*, for binding of a single ligand (*n* = 3 in the example). The graph vertices represent abstract conformations with patterns of ligand binding, denoted (*c_k_, S*), where *S* ⊆ {1, ···, *n*} is the subset of bound sites. Directed edges represent transitions arising either from binding without change of conformation (“vertical” edges), (*c_k_, S*) → (*c_k_, S* ∪ {*i*}) where *i* ∉ *S*, which occur for all conformations *c_k_*, or from conformational change without binding (“horizontal” edges), (*c_k_, S*) → (*c_j_, S*) where *k* ≠ *j*, which occur for all binding subsets *S*. Edges are shown in only one direction for clarity but are reversible, in accordance with thermodynamic equilibrium. Ignoring labels and thinking only in terms of vertices and edges, or “structure”, *A* has a product form: the vertical subgraphs, *A^c_k_^*, consisting of those vertices with conformation *c_k_* and all edges between them, all have the same structure and the horizontal subgraphs, *A_S_*, consisting of those vertices with binding subset *S* and all edges between them, also all have the same structure (Fig.3).

**Figure 3:**
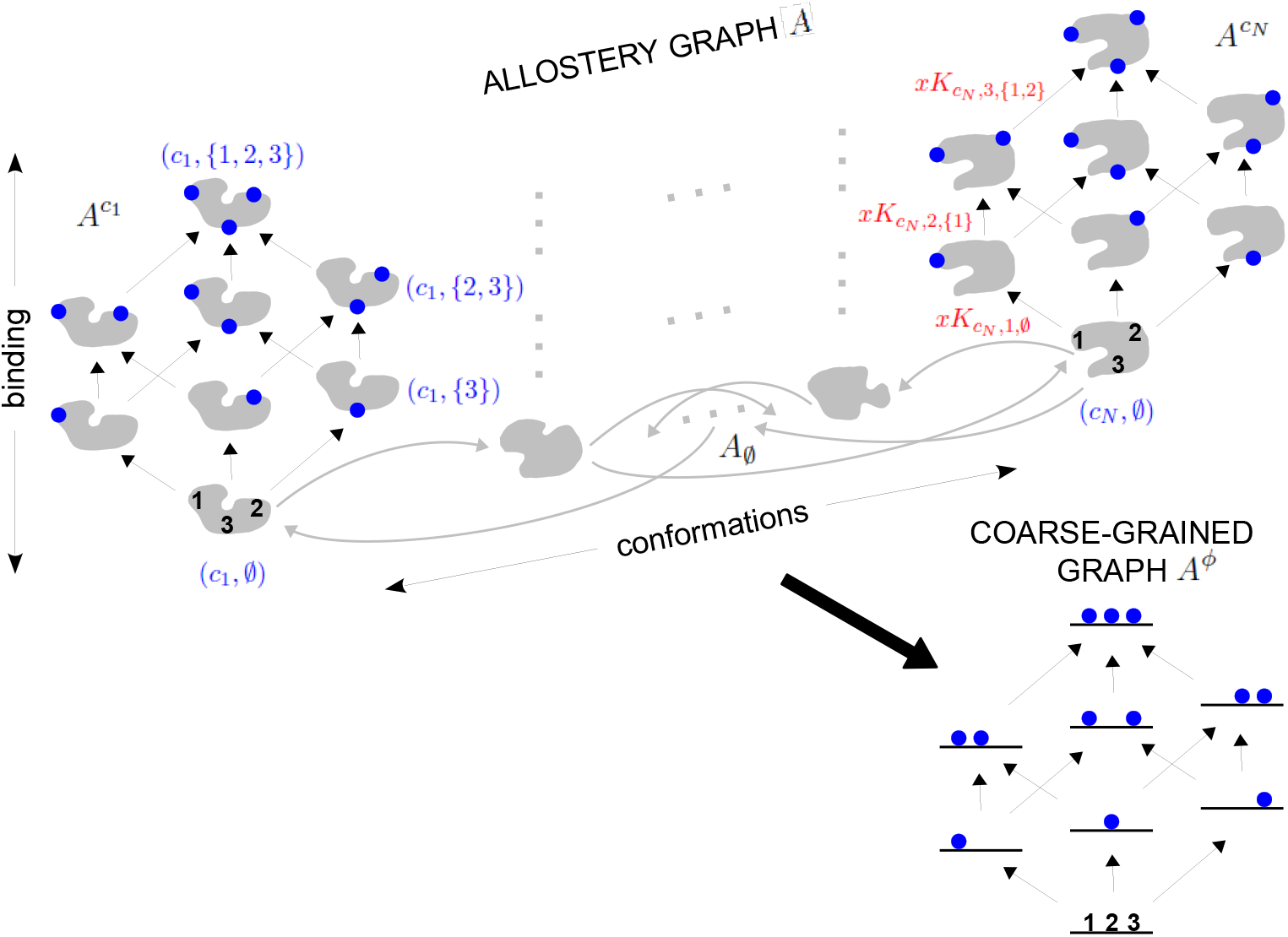
The allostery graph and coarse graining. A hypothetical allostery graph A (top) with three binding sites for a single ligand (blue discs) and conformations, *c*_1_, ···, *c_N_*, shown as distinct gray shapes. Binding edges (“vertical” in the text) are black and edges for conformational transitions (“horizontal”) are gray. Similar binding and conformational edges occur at each vertex but are suppressed for clarity. All vertical subgraphs, *A^c_k_^*, have the same structure, as seen for *A*^*c*_1_^ (left) and *A^c_N_^* (right), and all horizontal subgraphs, *A_S_*, also have the same structure, shown schematically for 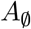 at the base. Example notation is given for vertices (blue font) and edge labels (red font), with *x* denoting ligand concentration and sites numbered as shown for vertices 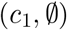 and 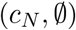. The coarse graining procedure coalesces each horizontal subgraph, *A_S_*, into a new vertex and yields the coarse-grained graph, *A^ϕ^* (bottom right), which has the same structure as *A^c_k_^* for any *k*. Further details in the text and the Materials and methods.

In an allostery graph, “conformation” is meant abstractly, as any state for which binding association constants can be defined. It does not imply any particular atomic configuration of a target molecule nor make any commitments as to how the pattern of binding changes.

The product-form structure of the allostery graph reflects the “conformational selection” viewpoint of MWC, in which conformations exist prior to ligand binding, rather than the “induced fit” viewpoint of KNF, in which binding can induce new conformations. Considerable evidence now exists for conformational selection, in the form of transient, rarely-populated conformations which exist prior to binding (Tzeng and Kalodimos, 2011). Induced fit may be incorporated within our graph-based approach by treating new conformations as always present but at extremely low probability. One of the original justifications for induced fit was that it enabled negative cooperativities, in contrast to conformational selection (Koshland and Hamadani, 2002), but we will show below that induced fit is not necessary for this and that negative HOCs arise naturally in our approach. Accordingly, the product-form structure of our allostery graphs is both convenient and powerful.

The edge labels are the non-dimensional ratios of the forward transition rate to the reverse transition rate; accordingly, the label for the reverse edge is the reciprocal of the label for the forward edge (Materials and methods). Labels may include the influence of components outside the graph, such as a binding ligand. For instance, the label for the binding edge (*c_k_, S*) → (*c_k_, S* ∪ {*i*}) is *xK_c_k_,i,S_*, where *x* is the ligand concentration and *K_c_k_,i,S_* is the association constant (Fig.1A), with dimensions of (concentration)^−1^, as described in the Introduction. Horizontal edge labels are not individually annotated and need only be specified for the horizontal subgraph of empty conformations, 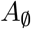, since all other labels are determined by detailed balance (Materials and methods).

The graph structure allows higher-order cooperativities (HOCs) between binding events to be calculated, as suggested in the Introduction. We will define this first for the “intrinsic” HOCs which arise in a given conformation and explain in the next section how “effective” HOCs are defined for the ensemble. In conformation *c_k_*, the intrinsic HOC for binding to site *i*, given that the sites in *S* are already bound, denoted *ω_c_k_,i,S_*, is defined by normalising the corresponding association constant to that for binding to site *i* when nothing else is bound, 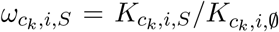 (Estrada et al., 2016). If *S* has only a single site, say *S* = {*j*}, then the HOC of order 1, *ω*_*c_k_,i*,{*j*}_, is the classical pairwise cooperativity between sites *i* and *j*. There is positive or negative HOC if *ω_c_k_,i,S_* > 1 or *ω_c_k_,i,S_* < 1, respectively, and independence if *ω_c_k_,i,S_* = 1 (Fig.1A).

For any graph *G*, the steady-state probabilities of the vertices can be calculated from the edge labels. For each vertex, *v*, in *G*, the probability, Pr_*v*_(*G*), is proportional to the quantity, *μ_v_*(*G*), obtained by multiplying the edge labels along any directed path of edges from a fixed reference vertex to *v*. It is a consequence of detailed balance that *μ_v_*(*G*) does not depend on the choice of path in *G*. This implies algebraic relationships among the edge labels. These can be fully determined from *G* and independent sets of parameters can be chosen (Materials and methods). For the allostery graph, a convenient choice vertically is those association constants *K_c_k_,i,S_* with *i* less than all the sites in *S*, denoted *i* < *S*; horizontal choices are discussed in the Materials and methods but are not needed for the main text.

Since probabilities must add up to 1, it follows that,

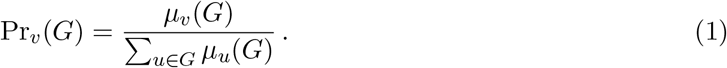

Eq.1 yields the same result as equilibrium statistical mechanics, with the denominator being the partition function for the thermodynamic grand canonical ensemble. HOCs provide a complete parameterisation at thermodynamic equilibrium. They replace the parameterisation by free energies of vertices, which is customary in physics, with a parameterisation by edge labels, at the expense of the algebraic dependencies described above. Equilibrium thermodynamics has also been used to define an overall measure of cooperativity for binding functions, for which higher-order free energies are introduced (Martini, 2017). These differ from our HOCs in also being associated to vertices rather than to edges. HOCs, higher-order free energies and conventional free energies offer alternative parameterisations of an equilibrium system which are readily inter-converted. Despite the lack of independence which arises from focussing on edges, HOCs are more effective for analysing information integration, precisely because they are associated to binding, which is the key mechanism through which information is conveyed to a target molecule.

Our specification of an allostery graph allows for arbitrary conformational complexity and arbitrary interacting ligands (we consider only one ligand here for simplicity), with the independent association constants in each conformation being arbitrary and with arbitrary changes in these parameters between conformations. Moreover, the abstract nature of “conformation”, as described above, permits substantial generality. Allostery graphs can be formulated to encompass the two conformations of MWC (Marzen et al., 2013), nested models (Robert et al., 1987), Cooper and Dryden’s fluctuations (Cooper and Dryden, 1984) and more recent views of dynamical allostery (Tzeng and Kalodimos, 2011), the multiple domains of the Ensemble Allosteric Model developed by Hilser and colleagues (Hilser et al., 2012) and applied also to intrinsically disordered proteins (Motlagh et al., 2012), other ensemble models (LeVine and Weinstein, 2015, Tsai and Nussinov, 2014) and Markov State Models arising from molecular dynamics simulations (Noé and Fischer, 2008).

### Coarse graining yields effective HOCs

As MWC showed, even if there is no intrinsic cooperativity in any conformation, an effective cooperativity can arise from the ensemble. This is usually detected in the shape of the binding function (Fig.2A). Here, we introduce a method of coarse-graining through which effective cooperativities can be rigorously defined. We illustrate this for the allostery graph, *A*, and explain the general coarse-graining method in the Materials and methods. For allostery, the idea is to treat the horizontal subgraphs, *A_S_*, as the vertices of a new coarse-grained graph, *A^ϕ^* (Fig.3, bottom right). There is an edge between two vertices in *A^ϕ^*, if, and only if, there is an edge in *A* between the corresponding horizontal subgraphs. It is not hard to see that *A^ϕ^* is identical in structure to any of the vertical subgraphs *A^c_k_^*. We can think of *A^ϕ^* as if it represents a single effective conformation to which ligand is binding and we can index each vertex of *A^ϕ^* by the corresponding subset of bound sites, *S*. The key point, as explained in the Materials and methods, is that it is possible to assign labels to the edges in *A^ϕ^* so that,

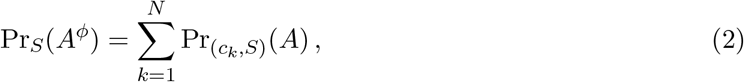

with *A^ϕ^* being at thermodynamic equilibrium under these label assignments. According to Eq.2, the probability of being in a coarse grained vertex of *A^ϕ^* is identical to the overall probability of being in any of the corresponding vertices of *A*. This is exactly the property a coarse graining should satisfy at steady state. This assignment of labels to *A^ϕ^* is the only way to ensure Eq.2 at equilibrium (Materials and methods), so that the coarse graining is both systematic and unique. It offers a general method for calculating how effective behaviour emerges, at thermodynamic equilibrium, from a more detailed underlying mechanism.

This method of coarse graining is likely to be broadly useful for other studies. We note that it may be considered as a way of calculating conditional probabilities and that it applies only to the steady state. It does not provide a coarse graining of the underlying dynamics, which is a much harder problem.

Because *A^ϕ^* resembles the graph for ligand binding at a single conformation, we can calculate HOCs for *A^ϕ^*—equivalently, effective HOCs for *A*—just as we did above, by normalising the effective association constants. Once the dust of calculation has settled (Materials and methods), we find that *A* has effective HOCs,

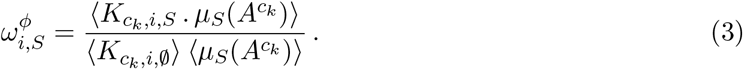

The quantity *μ_S_*(*A^c_k_^*) is calculated by multiplying labels over paths, as above, within the vertical subgraph *A^c_k_^*. The terms within angle brackets, of the form 〈*X*(*c_k_*)〉, where *X*(*c_k_*) is some function over conformations *c_k_*, denote averages over the steady-state probability distribution of the horizontal subgraph: 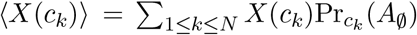. The formula in Eq.3 has a suggestive structure: it is an average of a product divided by the product of the averages. The resulting effective HOCs provide a biophysical language in which the integrative capabilities of any ensemble can be rigorously specified.

### Effective HOCs for MWC-like ensembles

The functional viewpoint is readily recovered from the ensemble. A generalised MWC formula can be given in terms of effective HOCs, from which the classical two-conformation MWC formula is easily derived (Materials and methods). Some expected properties of effective HOCs are also easily checked (Materials and methods). First, 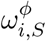 is independent of ligand concentration, *x*. Second, there is no effective HOC for binding to an empty conformation, so that 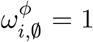. Third, if there is only one conformation *c*_1_, then the effective HOC reduces to the intrinsic HOC, so that 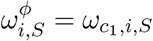.

More illuminating are the effective HOCs for the MWC model. We consider any conformational ensemble which is MWC-like: there is no intrinsic HOC in any conformation, so that *ω_c_k_,i,S_* = 1 and 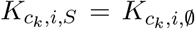; and the association constants are identical at all sites, so that we can set 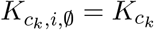. There may, however, be any number of conformations, not just the two conformations of the classical MWC model. It then follows that 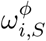 depends only on the size of *S*, so that we can write 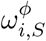 as 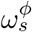, where *s* = #(*S*) is the order of cooperativity. Eq.3 then simplifies to (Materials and methods),

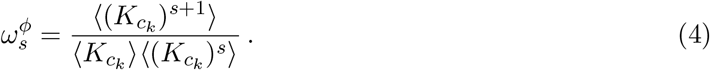

We see that, although there is no intrinsic HOC in any conformation, effective HOC of each order arises from the moments of *K_c_k__* over the probability distribution on 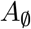. In particular, Eq.4 shows that the effective pairwise cooperativity is 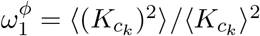.

In studies of G-protein coupled receptor (GPCR) allostery, Ehlert relates “empirical” to “ultimate” levels of explanation by a procedure similar to our coarse graining, but with only two conformations, and calculates a “cooperativity constant” which is the same as 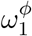 (Ehlert, 2016). Gruber and Horovitz calculate “successive ligand binding constants” for the two-conformation MWC model which are the same as effective association constants, 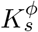, (Gruber and Horovitz, 2018) (Materials and methods). To our knowledge, these are the only other calculations of effective allosteric quantities. We note that Eq.4 applies to all HOCs, not just pairwise, and to any MWC-like ensemble, not just those with two conformations.

The classical MWC model yields only positive cooperativity (Koshland and Hamadani, 2002, Monod et al., 1965), as measured in the functional perspective (Fig.2A). We find that MWC-like ensembles yield positive effective HOCs of all orders. Strikingly, these effective HOCs increase with increasing order of cooperativity: provided *K_c_k__* is not constant over conformations (Materials and methods),

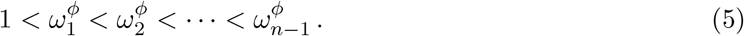

This shows that ensembles with independent and identical sites, including the two-conformation MWC model, can effectively implement high orders and high levels of positive cooperativity. Eq.5 is very informative and we return to it in the Discussion.

It is often suggested that negative cooperativity requires a different ensemble to those considered here, such as one allowing KNF-style induced fit (Koshland and Hamadani, 2002). However, if two sites are independent but not identical, so that 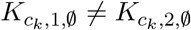, then, with just two conformations, the effective pairwise cooperativity can become negative. Indeed, 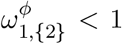, if, and only if, the values of the association constants are not in the same relative order in the two conformations (Materials and methods). Negative effective cooperativity requires distinct sites, not a distinct ensemble.

### Integrative flexibility of ensembles

Eq.3 shows that effective HOCs of any order can arise for a conformational ensemble but does not reveal what values they can attain. Can they vary arbitrarily? The question can be rigorously posed as follows. Suppose that numbers *α_i, S_* > 0 are chosen at will, with *i* < *S* to satisfy independence (above). Does there exist a conformational ensemble such that 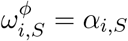?

To address this, we assume that there is no intrinsic HOC, so as not to introduce cryptically what we want to generate. It follows that the sites cannot be identical, for otherwise Eq.5 shows that integrative flexibility is impossible. Accordingly, the bare association constants, 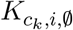 for 1 ≤ *i* ≤ *n*, can be treated as *n* free parameters in each conformation *c_k_*. If there are *N* conformations in the ensemble, then there are *N* – 1 free parameters coming from the horizontal edges (Materials and methods). Dimensional considerations imply that the effective HOCs cannot take arbitrary values if *n*(*N* – 1) < 2^*n*^ – 1. Conversely, but with more difficulty, we prove in the Materials and methods that any pattern of values can be realised, to any required degree of accuracy, provided there are enough conformations with the right probability distribution and patterns of association constants. Fig.4 illustrates this result. It shows three arbitrarily chosen patterns of positive and negative effective HOCs of all orders for a target molecule with four ligand binding sites and gives three corresponding conformational ensembles which exhibit those effective HOCs. We reiterate that this takes place without intrinsic cooperativity, so that binding remains independent in each individual conformation. Fig.4 also shows the complex binding behaviours exhibited by these ensembles, in terms of the average binding at each site (coloured curves), which can exhibit non-monotonicity because of negative effective HOCs. In contrast, the overall binding function (black curve) always increases.

**Figure 4:**
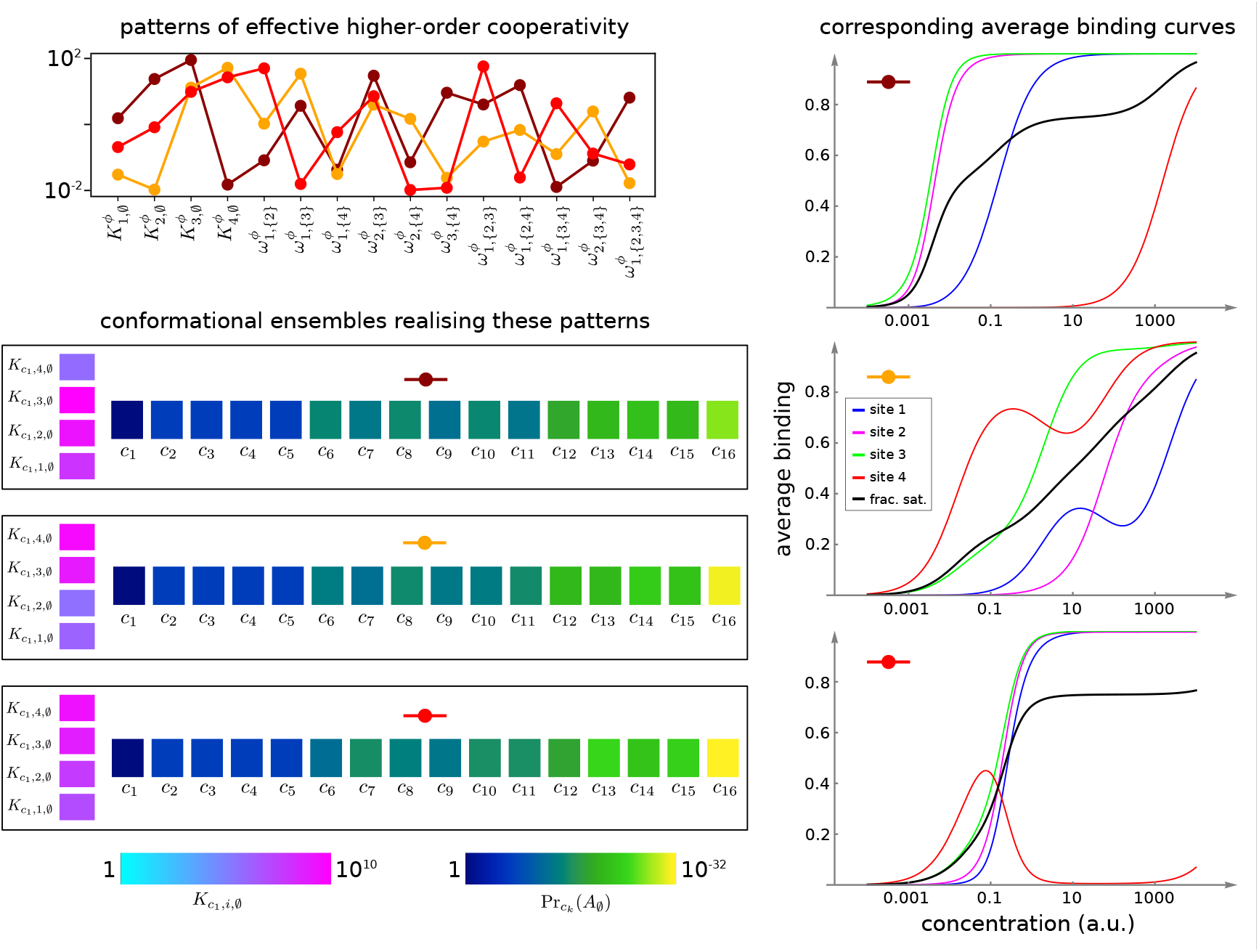
Integrative flexibility of allostery, illustrated for ligand binding to four sites. Top left panel shows three example patterns of effective bare association constants, 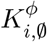, in arbitrary units of (concentration)^−1^, and effective HOCs, 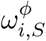 for *i* < *S*, in non-dimensional units; each example is coded by a colour (maroon, orange, red). Right panels show corresponding plots of average binding at each site (colours described in middle inset) and the binding function (black). Bottom left panels summarise the parameters of allostery graphs with *N* = 16 conformations, *c*_1_, ···, *c*_16_, exhibiting the effective parameters to an accuracy of 0.01 (Materials and methods), showing the intrinsic bare association constants in the reference conformation, *c*_1_ (left), and the probability distribution on the subgraph of empty conformations, 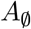 (right), colour coded as in the respective legends below. Intrinsic association constants for conformations other than *c*_1_ are determined in the proof (Materials and methods). Numerical values are given in the Materials and methods. Calculations were undertaken in a Mathematica notebook, available on request.

Our proof of integrative flexibility leaves open whether patterns of effective HOC can be reproduced exactly, rather than approximately, and with parameter values restricted to lie within specified ranges, but it confirms that there is no fundamental biophysical limitation to achieving any pattern of values. Accordingly, a central result of the present paper is that sufficiently complex allosteric ensembles can implement any form of information integration that is achievable without energy expenditure.

## DISCUSSION

Jacques Monod famously described allostery as “the second secret of life” (Ullmann, 2011). It is only relatively recently, however, that the prescience of his remark has been appreciated and the wealth of conformational ensembles present in most cellular processes has been revealed (Changeux and Christopoulos, 2016, Motlagh et al., 2014, Nussinov et al., 2013).

The present paper seeks to expand the existing allosteric perspective by providing a biophysical foundation for information integration by conformational ensembles. Eq.24 and Eq.25 in the Materials and methods (the latter being Eq.3 above) provide for the first time a rigorous definition of effective, higher-order quantities—the association constants, 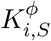, and cooperativities, 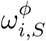—arising from any ensemble. Since our methods are equivalent to those of equilibrium statistical mechanics (Material and methods), these definitions correctly aggregate the free-energy contributions which emerge in the ensemble from ligand binding to a conformation, intrinsic cooperativity within a conformation and conformational change. As noted above, our results encompass recent work on effective properties of the classical, two-conformation MWC ensemble—for pairwise cooperativity (Ehlert, 2016) and higher-order association constants (Gruber and Horovitz, 2018)—but they hold more generally for ensembles of arbitrary complexity with any number of conformations, including those with intrinsic cooperativities.

The effective quantities introduced here provide a language in which the integrative capabilities of an ensemble can be rigorously expressed. To begin with, the overall binding function can be determined in terms of the effective quantities through a generalised MWC formula (Materials and methods), thereby recovering the functional viewpoint (Fig.2A) from the ensemble viewpoint (Fig.2B). This generalised MWC formula reduces to the usual MWC formula for the classical two-conformation MWC model (Eq.31). We also clarify issues which had been difficult to understand in the absence of a quantitative definition of effective quantities. We find that the classical MWC model exhibits effective HOCs of any order and that these are always positive. In other words, binding always encourages further binding. Moreover, these effective HOCs increase strictly with increasing order (Eq.5), so that the more sites which are bound, the greater the encouragement to further binding. We see that higher-order cooperativity has always been present, even for oxygen binding to haemoglobin, albeit unrecognised for lack of an appropriate quantitative definition. Eq.5 confirms in a more precise way the long-standing realisation from the functional perspective that the MWC model exhibits only positive cooperativity; at the same time it succinctly expresses the rigidity and limitations of this model.

It is often stated in the allostery literature that negative cooperativity requires induced fit, in which binding induces conformations which are not present prior to binding. This view goes back to Koshland, who pointed to the emergence of negative cooperativity in the KNF model of allostery, which allows induced fit, and contrasted that to the positive cooperativity of the MWC model, which assumes conformational selection (Koshland and Hamadani, 2002). Our language of effective quantities permits a more discriminating analysis. It confirms, as just pointed out, that the classical MWC model exhibits only positive effective HOCs but also shows that induced fit is not required for negative effective HOC, which can arise just as readily from conformational selection (Materials and methods). What is required is not a different ensemble but, rather, binding sites that are not identical.

Our main result, on the flexibility of conformational ensembles, shows that positive and negative HOCs of any value can occur in any pattern whatsoever, provided that the conformational ensemble is sufficiently complex, with enough conformations (Fig.4). Since the effective quantities provide a complete parameterisation of an ensemble at thermodynamic equilibrium, we see that conformational ensembles can implement any form of information integration that is achievable without external sources of energy.

The ability of conformational ensembles to flexibly implement information integration addresses a question raised in our previous work (Estrada et al., 2016), as to a molecular mechanism which can account for experimental data on gene regulation. Eukaryotic gene regulation is one of the most complex forms of cellular information processing (Wong and Gunawardena, 2020). Information from the binding of multiple transcription factors (TFs) at many sites, often widely distributed across the genome in distal enhancer sequences, must be integrated to determine whether, and in what manner, a gene is expressed. The results of the present paper offer a way to think further about how such integration takes place (Tsai and Nussinov, 2011). We focus on gene regulation but our results may also be useful for analysing other mechanisms of information integration, such as G-protein coupled receptors (Thal et al., 2018).

As pointed out in the Introduction, haemoglobin solves the problem of integrating two quite different physiological functions—picking up oxygen in the lungs and delivering oxygen to the tissues—by having two conformations, each adapted to one of these functions, and dynamically inter-converting between them (Fig.5A). The effective cooperativity of oxygen binding ensures that the appropriate conformation dominates the ensemble in the distinct contexts of the lungs, where oxygen is abundant, and the tissues, where oxygen is scarce, so that oxygen is transferred from the former to the latter.

**Figure 5:**
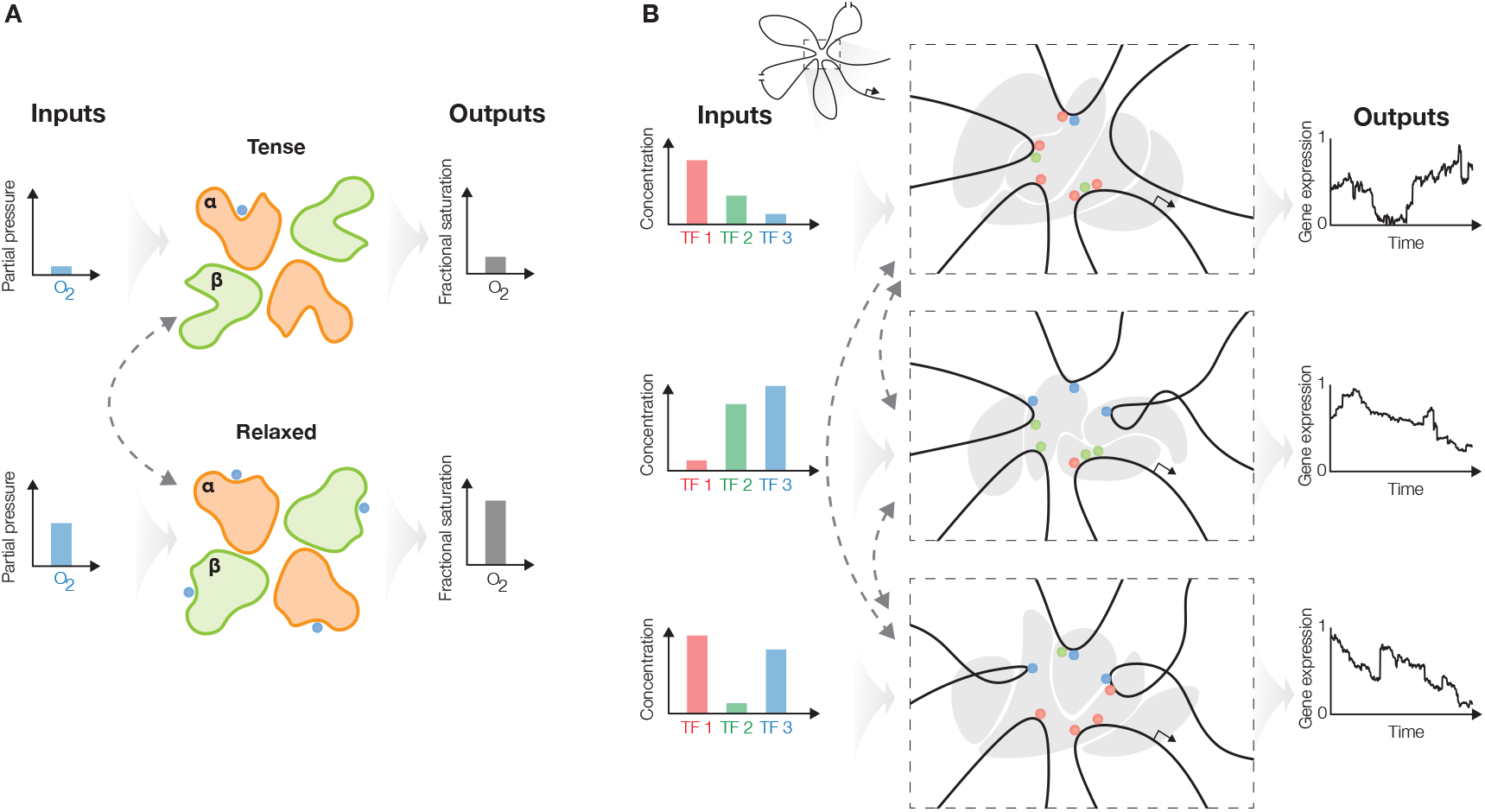
The haemoglobin analogy in gene regulation. **A.** The two conformations of haemoglobin are each adapted to one of the two input-output functions which haemoglobin integrates to solve the oxygen transport problem. These conformations dynamically interchange in the ensemble (grey dashed arrows). **B.** The gene regulatory machinery relates input patterns of TFs (left) to stochastic expression of mRNA (right). Our results suggest that a sufficiently complex conformational ensemble, built out of chromatin, TFs, co-regulators and phase-separated condensates (centre, gray shapes in three distinct conformations) could integrate these functions at a single gene in an analogous way to haemoglobin. Chromatin is represented by the thick black curve, whose looped arrangement around the promoter is shown schematically (top).

Genes have to be regulated to achieve yet more elaborate forms of integration, with the same gene being expressed differently in different contexts. Such pleiotropy is particularly evident in developmental genes (Bolt and Duboule, 2020) but usually occurs in distinct cells within the developing organism. The same gene is present in these cells but it may be difficult to know whether the corresponding regulatory machineries are also the same. More directly suitable examples for the present discussion arise in individual cells exposed to distinct stimuli (Molina et al., 2013, Kalo et al., 2015, Lin et al., 2015), which may be particularly the case for neurons or cells of the immune system (Marco et al., 2020, Smale et al., 2013).

Depending on the pattern of TFs present in a given cellular context (Fig.5B, left), a gene may be expressed in a certain way, as a distribution of splice isoforms, each with an overall level of mRNA expression and a pattern of stochastic bursting (Fig.5B, right). A different input pattern of TFs may elicit a different mRNA output. Our results suggest that one way in which these different input-output relationships could be integrated in the workings of a single gene is through allostery of the overall regulatory machinery. An allosteric analogy in gene regulation was previously made by Leonid Mirny, who suggested that the indirect cooperativity observed between transcription factors (TFs) could be mediated by nucleosomes (Mirny, 2010). In this view, TF binding to DNA takes place in one of two conformations—nucleosome present or absent—which dynamically inter-change, leading to the classical MWC model. Here, we build upon Mirny’s idea to suggest that not only indirect cooperativity but also, more broadly, information integration may be accounted for by the conformational dynamics of the gene regulatory machinery. The latter comprises not just individual nucleosomes but whatever other molecular entities are implicated in conveying information from TF binding sites to RNA polymerase and the transcriptional machinery (Fig.5B, centre), as discussed below. If this hypothesis is correct then the flexibility result tells us that the overall regulatory conformational ensemble must exhibit substantial complexity to implement genetic integration.

Studies of individual regulatory components have revealed many levels of conformational complexity. DNA itself exhibits conformational changes in respect of TF binding (Kim et al., 2013). Nucleosomes are moved or evicted to alter chromatin conformation and DNA accessibility (Mirny, 2010, Voss and Hager, 2014). TFs, in particular, show high levels of intrinsic disorder compared to other classes of proteins (Liu et al., 2006), especially in their activation domains, and these disordered regions exhibit dynamic multivalent interactions characteristic of higher-order effects (Chong et al., 2018, Clark et al., 2018). Hub TFs like p53 exhibit high levels of conformational flexibility in the context of specific DNA binding (Demir et al., 2017). Transcriptional co-regulators, which do not directly bind DNA but are recruited there by TFs, exhibit substantial conformational complexity: CBP/p300 has multiple intrinsically disordered regions which facilitate higher-order cooperative interactions (Dyson and Wright, 2016), while the Mediator complex exhibits quite remarkable conformational changes upon binding to TFs (Allen and Taatjes, 2015). Transcription initiation sub-complexes such as TFIID, which help assemble the transcriptional machinery, show conformational plasticity (Nogales et al., 2017), while the C-terminal domain of RNA Pol II, which is repetitive and intrinsically disordered, shows surprising local structural heterogeneity (Portz et al., 2017). The significance of RNA conformational dynamics during transcription is becoming clearer (Ganser et al., 2019). Finally, transcription may also be regulated within larger-scale entities, such as transcription factories (Edelman and Fraser, 2012), phase-separated condensates (Sabari et al., 2018) and topological domains (Benabdallah and Bickmore, 2015). The role of such entities remains a matter of debate (Mir et al., 2019) but they may play a significant role in conveying information over long genomic distances between distal enhancers and target promoters (Furlong and Levine, 2018). From the perspective taken here, in view of their size and extent, they may exhibit conformational dynamics on longer timescales.

These various findings have emerged largely independently of each other. They indicate the presence of many conformations of components of the gene regulatory machinery, with these components dynamically interchanging on varying timescales. The collective effect of these coupled dynamics is difficult to predict but we can hazard some guesses. It has been suggested, for example, that multi-protein complexes like Mediator couple the conformational repertoires of their components proteins into complex allosteric networks for processing information (Lewis, 2010). From an ensemble viewpoint, if component *X* has *m* conformations and component *Y* has *n* conformations, we might naively expect that the coupling of *X* and *Y* in a complex yields roughly *mn* conformations. Following this multiplicative logic for the many components involved in eukaryotic gene regulation, from DNA itself to condensates and domains, suggests that the gene regulatory machinery has enormous conformational capacity with a deep hierarchy of timescales.

In making the analogy to haemoglobin, it is the conformational dynamics which implements the transfer of information from upstream TF inputs to downstream gene output. In any given cellular context, as determined by the input pattern of TFs, we may expect one, or perhaps a few, overall regulatory conformations which are well-adapted to generate the required mRNA output and these conformations will be the most frequently observed. The ensemble may exhibit complex patterns of positive and negative effective HOCs among the input TFs which will characterise the required output. In the light of our flexibility theorem, the occurrence of such HOCs, which appear to be necessary to account for data on gene regulation (Park et al., 2019a), may be seen as evidence for conformational complexity. When the cellular context changes, different conformations, adapted to produce the output required in the new context, may be present most often—although careful inspection may show them to have been more fleetingly present previously, as would be expected under conformational selection. More broadly, the complexity of the regulatory conformational ensemble and its dynamics reflect the complexity of functional integration which the gene has to undertake.

Furlong and Levine have suggested a “hub and condensate” model for the overall gene regulatory machinery, which brings together aspects of earlier models to account for how remote enhancers communicate with target promoters (Furlong and Levine, 2018). The allosteric perspective taken here emphasises the significance of conformational dynamics for the functional integration under-taken by such “hubs”.

Testing these ideas on the scale of the regulatory machinery presents a daunting challenge but recent developments point the way towards approaching them, including advances in cryo-EM (Lewis and Costa, 2020), single-molecule microscopy (Li et al., 2019, Bacic et al., 2020), NMR (Shi et al., 2020), synthetic biology (Park et al., 2019b) and the measurement of higher-order quantities (Gruber and Horovitz, 2018). Before experiments can be formulated, an appropriate conceptual picture needs to be described and that is what we have tried to formulate here. We now know a great deal about the molecular components involved in gene regulation but the question of how these components collectively give rise to function has been harder to grasp. The allosteric analogy to haemoglobin, upon which we have built here, suggests a potential way to fill this gap.

In extending the haemoglobin analogy, we have sidestepped the issue of energy expenditure. This is not relevant for haemoglobin but it can hardly be avoided in considering eukaryotic gene regulation, where reorganisation of chromatin and nucleosomes requires energy-dissipating motor proteins and post-translational modifications driven by chemical potential differences are found on all components of the regulatory machinery. What impact such energy expenditure has on ensemble functional integration is a very interesting question. In the light of what is currently known (Estrada et al., 2016, Biddle et al., 2019, Wong and Gunawardena, 2020), we expect that energy expenditure will improve the functional capabilities of an allosteric ensemble, beyond what can be achieved at thermodynamic equilibrium. For this reason, we feel the discussion presented here still offers a reasonable starting point for thinking about how regulatory ensembles integrate genetic information. If, indeed, regulatory energy expenditure is essential for gene expression function, as experimental evidence increasingly suggests (Park et al., 2019a), new methods, both theoretical and experimental, will be required to understand its functional significance.

## MATERIAL AND METHODS

### THE LINEAR FRAMEWORK

#### Background and references

The graphs described in the main text, like those in Fig.3, are “equilibrium graphs”, which are convenient for describing systems at thermodynamic equilibrium. Equilibrium graphs are derived from linear framework graphs. The distinction between them is that the latter specifies a dynamics, while the former specifies an equilibrium steady state. We first explain the latter and then describe the former. Throughout this section we will use “graph” to mean “linear framework graph” and “equilibrium graph” to mean the kind of graph used in the main text.

The linear framework was introduced in (Gunawardena, 2012), developed in (Mirzaev and Gunawardena, 2013, Mirzaev and Bortz, 2015), applied to various biological problems in (Ahsendorf et al., 2014, Dasgupta et al., 2014, Estrada et al., 2016, Wong et al., 2018a,b, Yordanov and Stelling, 2018, Biddle et al., 2019, Yordanov and Stelling, 2020) and reviewed in (Gunawardena, 2014, Wong and Gunawardena, 2020). Technical details and proofs of the ideas described here can be found in (Gunawardena, 2012, Mirzaev and Gunawardena, 2013) as well as in the Supplementary Information of (Estrada et al., 2016, Wong et al., 2018b, Biddle et al., 2019).

The framework uses finite, directed graphs with labelled edges and no self-loops to analyse biochemical systems under timescale separation. In a typical timescale separation, the vertices represent “fast” components or states, which are assumed to reach steady state; the edges represent reactions or transitions; and the edge labels represent rates with dimensions of (time)^−1^. The labels may include contributions from “slow” components, which are not represented by vertices but which interact with them, such as binding ligands in the case of allostery.

#### Linear framework graphs and dynamics

Graphs will always be connected, so that they cannot be separated into sub-graphs between which there are no edges. The set of vertices of a graph *G* will be denoted by *ν*(*G*). For a general graph, the vertices will be indexed by numbers 1, ⋯, *N* ∈ *ν* (*G*) and vertex 1 will be taken to be the reference vertex. Particular kinds of graphs, such as the allostery graphs discussed in the Paper, may use a different indexing. An edge from vertex *i* to vertex *j* will be denoted *i* → *j* and the label on that edge by ℓ(*i* → *j*). A subscript, as in *i* →*_G_ j*, may be used to specify which graph is under discussion. When discussing graphs, we used the word “structure” to refer to properties that depend on vertices and edges only, ignoring the labels.

A graph gives rise to a dynamical system by assuming that each edge is a chemical reaction under mass-action kinetics with the label as the rate constant. Since each edge has only a single source vertex, the corresponding dynamics is linear and can be represented by a linear differential equation in matrix form,

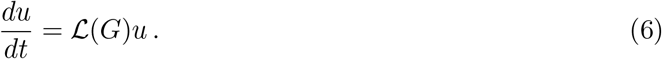

Here, *G* is the graph, *u* is a vector of component concentrations and 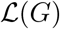 is the Laplacian matrix of *G*. Since material is only moved between vertices, there is a conservation law, 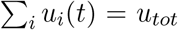. By setting *u_tot_* = 1, *u* can be treated as a vector of probabilities. In such a stochastic setting, Eq.6 is the master equation (Kolmogorov forward equation) of the underlying Markov process. This is a general representation: given any well-behaved Markov process on a finite state space, there is a graph, whose vertices are the states, for which Eq.6 is the master equation.

The linear dynamics in Eq.6 gives the linear framework its name and is common to all applications. The treatment of the external components, which appear in the edge labels and which introduce nonlinearities, depends on the application. For the case of allostery treated here, we assume that each ligand is present in a conserved total amount and that binding and unbinding to graph vertices do not change the free concentrations of the ligands. In this case, the edge labels are effectively constant. The same assumptions are implicitly used in other studies of allostery and reflect, from an equilibrium thermodynamic perspective, the assumptions of the grand canonical ensemble.

#### Steady states and thermodynamic equilibrium

The dynamics in Eq.6 always tends to a steady state, at which *du*/*dt* = 0, and, under the fundamental timescale separation, it is assumed to have reached a steady state. If the graph is strongly connected, it has a unique steady state up to a scalar multiple, so that dim ker 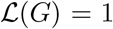. Strong connectivity means that, given any two distinct vertices, *i* and *j*, there is a path of directed edges from *i* to *j*, *i* = *i*_1_ → *i*_2_ → ⋯ → *i*_*k*−1_ → *i_k_* = *j*. Under strong connectivity, a representative steady state for the dynamics, 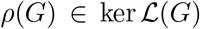, may be calculated in terms of the edge labels by the Matrix Tree Theorem. We omit the corresponding expression, as it is not needed here, but it can be found in any of the references given above. This expression holds whether or not the steady state is one of thermodynamic equilibrium. However, at thermodynamic equilibrium, the description of the steady state simplifies considerably because detailed balance holds. This means that the graph is reversible, so that, if *i* → *j*, then also *j* → *i*, and each pair of such edges is independently in flux balance, so that,

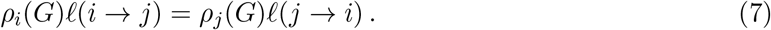

This “microscopic reversibility” is a fundamental property of thermodynamic equilibrium. Note that a reversible, connected graph is necessarily strongly connected.

Take any path of reversible edges from the reference vertex 1 to some vertex *i*, 1 = *i*_1_ ⇌ *i*_2_ ⇌ ⇌ *i*_*k*−1_ → *i_k_* = *i*, and let *μ_i_*(*G*) be the product of the label ratios along the path,

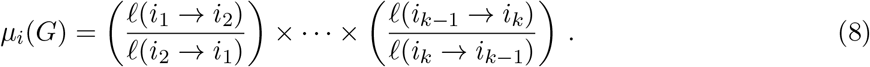

It is straightforward to see from Eq.7 that *μ_i_*(*G*) does not depend on the chosen path and that *ρ_i_*(*G*) = *μ_i_*(*G*)*ρ*_1_ (*G*). The vector *μ*(*G*) is therefore a scalar multiple of *ρ*(*G*) and so also a steady state for the dynamics. The detailed balance formula in Eq.7 also holds for *μ* in place of *ρ*. At thermodynamic equilibrium, the only parameters needed to describe steady states are label ratios.

#### Equilibrium graphs and independent parameters

This observation about label ratios leads to the concept of an equilibrium graph. Suppose that *G* is a linear framework graph which can reach thermodynamic equilibrium and is therefore reversible (above). *G* gives rise to an equilibrium graph, 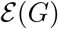 as follows. The vertices and edges of 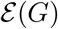 are the same as those of *G* but the edge labels in 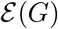, which we will refer to as “equilibrium edge labels” and denote ℓ_eq_(*i* → *j*), are the label ratios in *G*. In other words,

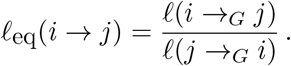

Note that the equilibrium edge labels of 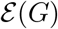 are non-dimensional and that ℓ_eq_(*j* → *i*) = ℓ_eq_(*i* → *j*)^−1^. The equilibrium edge labels are the essential parameters for describing a state of thermodynamic equilibrium.

These parameters are not independent because Eq.7 implies algebraic relationships among them. Indeed, Eq.7 is equivalent to the following “cycle condition”, which we formulate for 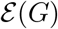: given any cycle of edges, *i*_1_ → *i*_2_ → ⋯ → *i*_*k*–1_ → *i*_1_, the product of the equilibrium edge labels along the cycle is always 1,

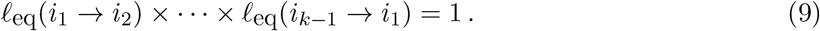

This cycle condition is equivalent to the detailed balance condition in Eq.7 and either condition is equivalent to *G* being at thermodynamic equilibrium.

There is a systematic procedure for choosing a set of equilibrium edge label parameters which are both independent, so that there are no algebraic relationships among them, and also complete, so that all other equilibrium edge labels can be algebraically calculated from them. Recall that a spanning tree of *G* is a connected subgraph, *T*, which contains each vertex of *G* (spanning) and which has no cycles when edge directions are ignored (tree). Any strongly connected graph has a spanning tree and the number of edges in such a tree is one less than the number of vertices in the graph. Since *G* and 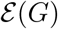 have the same vertices and edges, they have identical spanning trees. The equilibrium edge labels ℓ_eq_(*i* →*_T_ j*), taken over all edges *i* → *j* of *T*, form a complete and independent set of parameters at thermodynamic equilibrium. In particular, if *G* has *N* vertices, there are *N* – 1 independent parameters at thermodynamic equilibrium.

In the main text, we defined an equilibrium allostery graph, *A* (Fig.3), without specifying a corresponding linear framework graph, *G*, for which 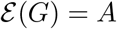. Because label ratios are used in an equilibrium graph, there is no unique linear framework graph corresponding to it. However, some choice of transition rates, ℓ(*i* →*_G_ j*) and ℓ(*j* →*_G_ i*), can always be made such that their ratio is 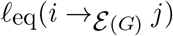. Hence, some linear framework graph *G* can always be defined such that 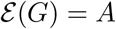. In some of the constructions below, we will work with the linear framework graph, *G*, rather than with the equilibrium graph *A* and will then show that the construction does not depend on the choice of *G*.

#### Steady-state probabilities and equilibrium statistical mechanics

The steady-state probability of vertex *i*, Pr*_i_*(*G*), can be calculated from the steady state of the dynamics by normalising, so that,

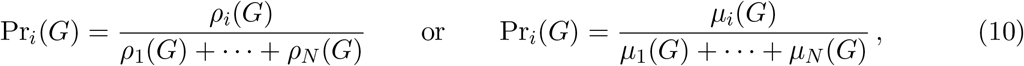

where the first formula holds for any strongly-connected graph and the second formula also holds if the graph is at thermodynamic equilibrium. In the latter case, Eq.7 holds and *μ*(*G*) can be defined by Eq.8. The second formula in Eq.10 corresponds to Eq.1. If the graph is at thermodynamic equilibrium, the equilibrium edge labels may be interpreted thermodynamically,

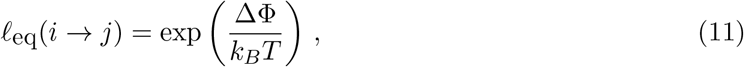

where ΔΦ is the free energy difference between vertex *i* and vertex *j*, *k_B_* is Boltzmann’s constant and *T* is the absolute temperature. If Eq.11 is used to expand the second formula in Eq.10, it gives the specification of equilibrium statistical mechanics for the grand canonical ensemble, with the denominator being the partition function.

It will be helpful to let ∏(*G*) and Ψ(*G*) denote the corresponding denominators in Eq.10, so that, Π(*G*) = *ρ*_1_(*G*) + ⋯ + *ρ_N_*(*G*) for any strongly-connected graph and Ψ(*G*) = *μ*_1_(*G*) + ⋯ + *μ_N_*(*G*) for a graph which is at thermodynamic equilibrium. We will refer to ∏(*G*) and Ψ(*G*) as partition functions. It follows from Eq.10 that,

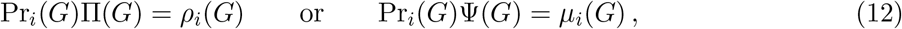

depending on the context.

### THE ALLOSTERY GRAPH

#### Structure and labels

An allostery graph, *A*, is an equilibrium graph which describes the interplay between conformational change and ligand binding. As defined in the main text, its vertices are indexed by (*c_k_, S*), where *c_k_* specifies a conformation with 1 ≤ *k* ≤ *N* and *S* ⊆ {1, ⋯, *n*} specifies a subset of sites bound by a ligand whose concentration is *x*. There is no difficulty in allowing multiple ligands and overlapping binding sites but to keep the formalism simple, we describe here the case of a single ligand.

Recall from the main text that *A* has vertical subgraphs, *A^c_k_^*, consisting of vertices (*c_k_, R*) for all binding subsets *R*, together with all edges between them, with the vertices indexed by binding subsets, *R*, and with 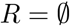 being the reference vertex. *A* has horizontal subgraphs, *A_S_*, consisting of vertices (*c_i_, S*) for all conformations *c_i_*, together with all edges between them, with the vertices labelled by conformations *c_i_*, and with *c*_1_ being the reference vertex. The product structure of *A* is revealed by all vertical subgraphs having the same structure as each other and all horizontal subgraphs having the same structure as each other (Fig.3). Scheme 1 below illustrates this further.

**Scheme 1:**
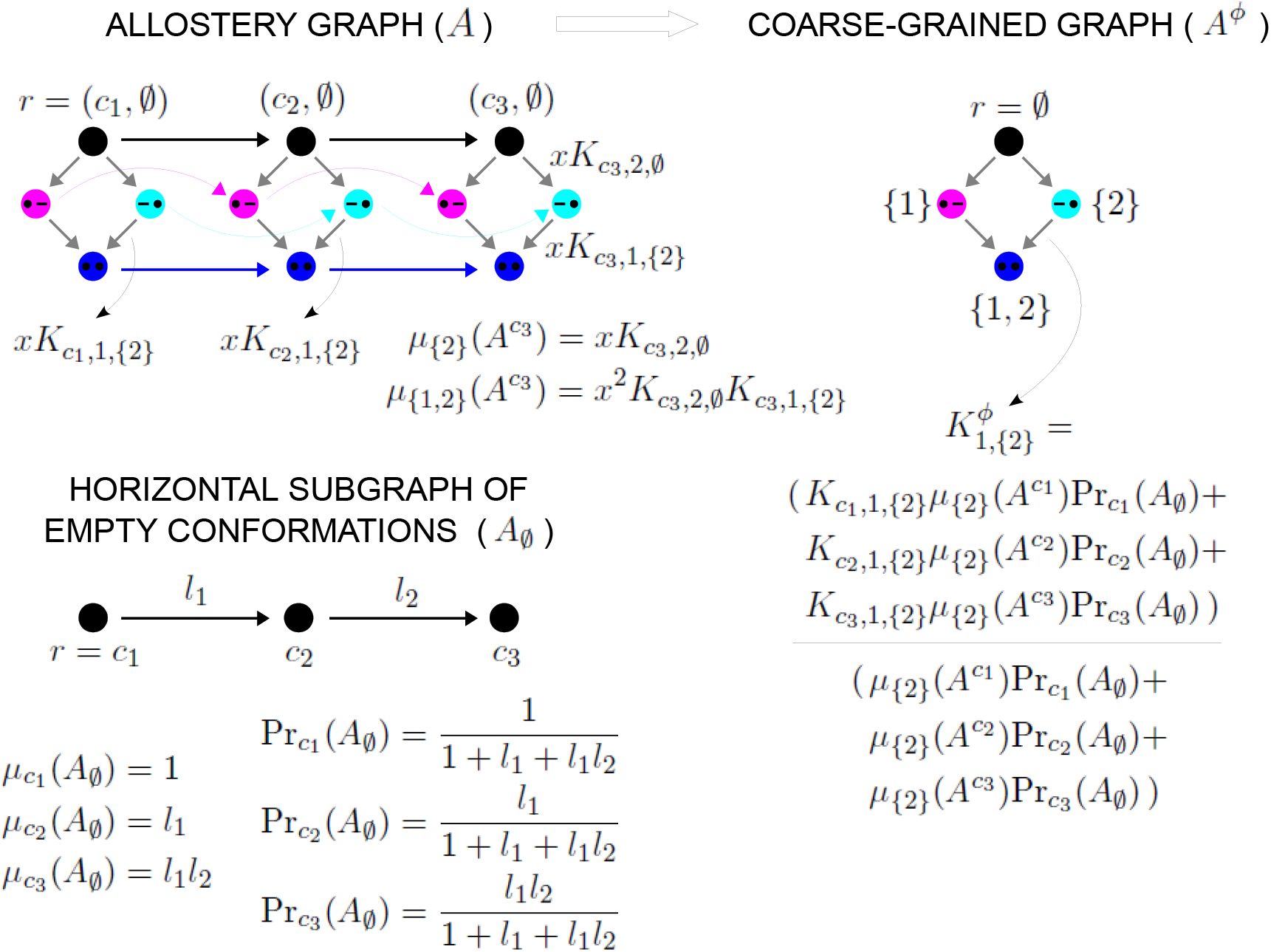
Coarse graining and effective association constants. At top left is an example allostery graph, with binding of a single ligand to *n* = 2 sites for *N* = 3 conformations. Vertices indicate a bound site with a solid black dot and an unbound site with a black dash and binding subsets are colour coded: both sites unbound, black; only site 1 bound, magenta; only site 2 bound, cyan; both sites bound, blue. Some vertices are annotated and some edge labels are shown, with x denoting ligand concentration. Example calculations of *μ_S_* based on Eq.8 are shown for the vertical subgraph *A*^*c*_3_^. At bottom is the horizontal subgraph 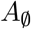 along with the calculation of its steady-state probability distribution in terms of the equilibrium labels, *l*_1_, *l*_2_ and the quantities *μ_c_k__*. At top right is the coarse-grained allostery graph, *A^ϕ^*, with vertices colour coded as for the binding subsets of the allostery graph. Eq.24 for the effective association constants is illustrated below *A^ϕ^*.

As for the labels, the vertical binding edges have equilibrium labels,

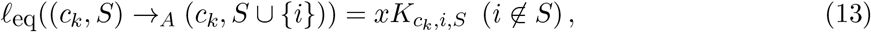

where *x* is the concentration of the ligand and *K_c_k_,i,S_* is the association constant for binding to site *i* when the ligand is already bound at the sites in *S*. The horizontal edges, which represent transitions between conformations, have equilibrium labels, ℓ_eq_((*c_k_, S*) →*_A_* (*c_l_, S*)), which are not individually annotated. However, it is only necessary to specify these equilibrium labels for a single horizontal subgraph, of which the subgraph of empty conformations, 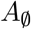, is particularly convenient. To see this, let us calculate the steady-state, *μ*_(*c_k_,S*)_(*A*), using Eq.8. Taking the reference vertex in *A* to be 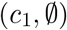, we can always find a path to any given vertex (*c_k_, S*) of *A*, by first moving horizontally within 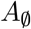 from 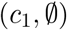 to 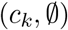 and then moving vertically within *A^c_k_^* from 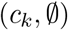 to (*c_k_, S*). According to Eq.8, the steady state is given by the product of the equilibrium labels along this path, so that,

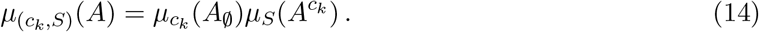

Now consider any horizontal edge in *A*, (*c_k_, S*) → (*c_l_, S*). Since *A* is at thermodynamic equilibrium, it follows from Eq.7, using *μ* in place of *ρ*, and Eq.14 that,

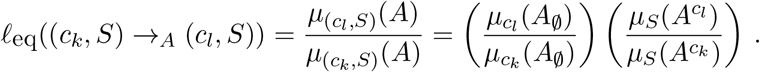

Applying Eq.7 to 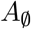, with *μ* in place of *ρ*, we see that

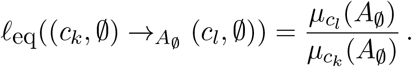

Hence, it follows that,

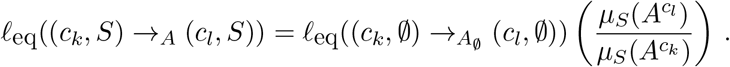

Accordingly, all the labels in *A* are determined by the vertical labels in Eq.13, from which *μ_S_*(*A^c_k_^*) and *μ_S_*(*A^c_l_^*) are determined, and the horizontal labels in the subgraph of empty conformations, 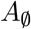.

#### Independent parameters

We can choose any spanning tree in the horizontal subgraph of empty conformations, 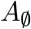. As explained above, the equilibrium labels on the edges of this tree define a complete set of *N* – 1 independent parameters for 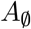. As for the vertical subgraphs, *A^c_k_^*, which all have the same structure, consider the subgraph of *A^c_k_^* consisting of all edges, together with the corresponding source and target vertices, of the form, (*c_k_, S*) → (*c_k_, S* ∪ {*i*}), where 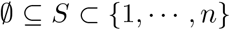 and *i* is less than all the sites in *S* (*i* < *S*). It is not difficult to see that this subgraph is a spanning tree of *A^c_k_^* (Estrada et al., 2016, SI, §3.2). Accordingly, the association constants, *K_c_k_,i,S_* from Eq.13, with *i* < *S*, form a complete set of independent parameters for *A^c_k_^*. Because of the product structure of *A*, adjoining the spanning trees in *A^c_k_^*, for each conformation *c_k_* with 1 ≤ *k* ≤ *N*, to the spanning tree in 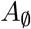, yields a spanning tree in *A*. Hence, the independent parameters for *A^c_k_^* together with the *N* – 1 independent parameters for 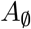 are also collectively independent as parameters for *A*. It follows from the description of labels above that these parameters are also complete for *A*, so that any equilibrium label in *A* can be expressed in terms of them.

### A GENERAL METHOD OF COARSE GRAINING

#### Coarse graining a linear framework graph and Eq.2

We will describe the coarse-graining procedure for an arbitrary reversible linear framework graph, *G*, and then explain how this can be adapted to an equilibrium graph, as described for the allostery graph *A* in the main text.

We will say that a graph *G* is *in-uniform* if, given any vertex *j* ∈ *v*(*G*), then for all edges *i* → *j*, ℓ(*i* → *j*) does not depend on the source vertex *i*.

##### Lemma 1

*Suppose that G is reversible and in-uniform. Then, G is at thermodynamic equilibrium and the vector θ given by θ_j_ = ℓ(i → j), which is well-defined by hypothesis, is a basis element in* 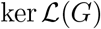 *and a steady state for the dynamics*.

**Proof:** If *i*_1_ ⇌ *i*_2_ ⇌ ⋯ ⇌ *i*_*i*−1_ ⇌ *i_k_* is any path of reversible edges in *G*, then the product of the label ratios along the path satisfies,

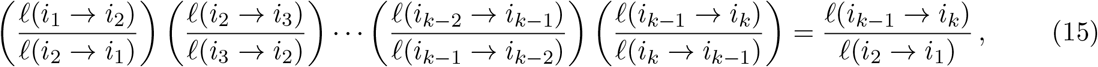

because the intermediate terms cancel out by the in-uniform hypothesis. If the path is a cycle, so that *i_k_* = *i*_1_, then, again because of the in-uniform hypothesis, the right-hand side of Eq. 15 is 1. Hence, *G* satisfies the cycle condition in Eq.9 and is therefore at thermodynamic equilibrium. For the last statement, assume that *i*_1_ is the reference vertex 1 and that *i_k_* = *j*, for any vertex *j*. Using Eq.8, we see that *μ_j_* (*G*) = *θ_j_*/*θ*_1_. Since *θ*_1_ is a scalar multiple, the last statement follows.

Now let *G* be an arbitrary reversible graph, which need not satisfy detailed balance. Let *G*_1_, ⋯, *G_m_* be any partition of the vertices of *G*, so that *G_i_* ⊆ *v*(*G*), *G*_1_ ∪ ⋯ ∪ *G_m_* = *v*(*G*) and 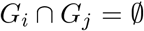 when *i* ≠ *j*. Let 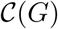 be the labelled directed graph with 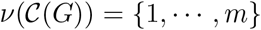 and let 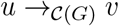 if, and only if, there exists *i* ∈ *G_u_* and *j* ∈ *G_v_* such that *i* →*_G_ j*. Finally, let the edge labels of 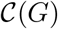 be given by,

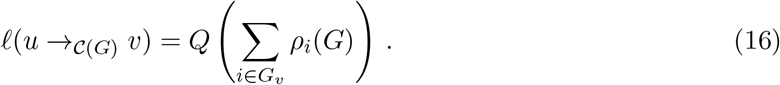

The quantity *Q* in Eq.16 is chosen arbitrarily so that the dimension of ℓ(*u* → *v*) is (time)^−1^, as required for an edge label. This is necessary because, by the Matrix Tree Theorem, the dimension of *ρ_i_*(*G*) is (time)^1−*N*^, where *N* is the number of vertices in *G*. However, *Q* plays no role in the analysis which follows because the coarse graining applies only to the steady state of 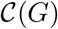, not its transient dynamics, and, as we will see, 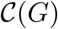 is always at thermodynamic equilibrium, so that *Q* disappears when equilibrium edge labels are considered.

Note that 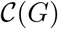 inherits reversibility from *G* and that 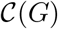 is in-uniform. Hence, by Lemma 1, 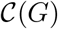 is at thermodynamic equilibrium and,

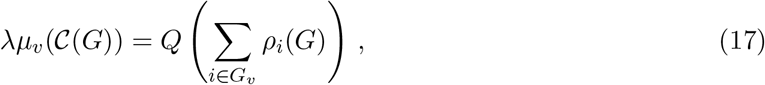

where λ is a scalar that does not depend on 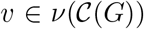. Since *G*_1_, ⋯, *G_m_* is a partition of the vertices of *G*, it follows from Eq.17 that

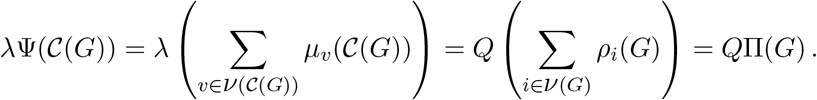

Eqs.12 and 17 then show that both λ and *Q* cancel in the ratio for the steady-state probabilities, so that,

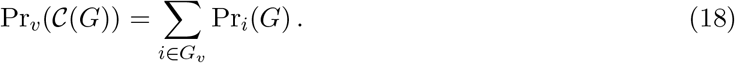

Eq.18 is the coarse-graining equation, as given in Eq.2.

#### Coarse graining an equilibrium graph

The coarse graining procedure described above can be applied to any reversible graph, which need not be at thermodynamic equilibrium. However, the coarse-graining described in the Paper was for an equilibrium graph. It is not difficult to see that the construction above can be undertaken consistently for any equilibrium graph. It is helpful to first establish a more general observation. The choice of edge labels for 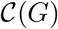, as given in Eq.16, is not the only one for which Eq.18 holds, as the appearance of the factor *Q* indicates. However, the label ratios in 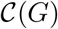 are uniquely determined by the labels of *G*.

Suppose that *G* is a reversible graph with a vertex partition *G*_1_, ⋯, *G_m_*, as above. *G* need not be at thermodynamic equilibrium. Suppose that *C* is a graph which is isomorphic to 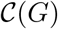 as a directed graph (“structurally isomorphic”), in the sense that it has identical vertices and edges but may have different edge labels. (Technically speaking, an “isomorphism” allows for the vertices of *C* to have an alternative indexing to those of 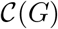, as long as the two indexings can be interconverted so as to preserve the edges. For simplicity of exposition, we assume that the indexing is, in fact, identical. No loss of generality arises from doing this.)

##### Lemma 2

*Suppose that C is at thermodynamic equilibrium and the coarse-graining equation (Eq.18) holds for C*, *so that* 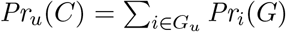. *If u ⇌_C_ v is any reversible edge, then its equilibrium label depends only on G*,

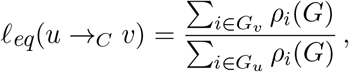

*and C and* 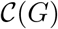 *are isomorphic as equilibrium graphs, so that identical edges have identical equilibrium labels*.

**Proof:** It follows from Eq.12 that Pr*_i_*(*G*) = *ρ_i_*(*G*)/Π(*G*) and, since *C* is at thermodynamic equilibrium, Pr*_u_*(*C*) = *μ_u_*(*C*)/Ψ(*C*). Using the coarse-graining equation for Pr*_u_*(*C*), we see that

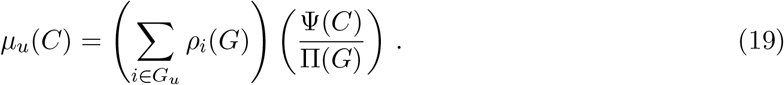

Since *C* is at thermodynamic equilibrium, Eq.7, with *μ* in place of *ρ*, implies that,

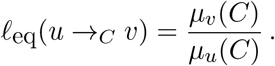

Substituting with Eq.19, the partition functions cancel out to give the formula above. Since 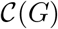 satisfies the same assumptions as *C*, it has the same equilibrium labels. Hence, *C* and 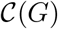 must be isomorphic as equilibrium graphs.

##### Corollary 1

*Suppose that A is an equilibrium graph and that G is any graph for which* 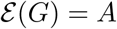, *as described above. If any coarse graining of G is undertaken to yield the coarse-grained graph* 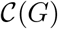, *which must be at thermodynamic equilibrium, then*

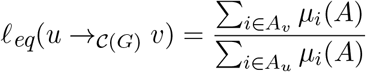

*and* 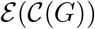 *depends only on A and not on the choice of G*.

**Proof.** *A* acquires from *G* the same coarse graining, with the partition *A*_1_, ⋯, *A_m_* of *v*(*A*), where *A_i_* = *G_i_* ⊆ {1, ⋯ *m*}. By hypothesis, *G* is at thermodynamic equilibrium, so that *ρ_i_*(*G*) = λ*μ_i_*(*G*) for some scalar multiple λ. Also, since 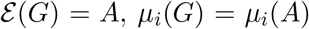. Substituting in the formula in Lemma 2 yields the formula above. The equilibrium labels of 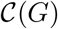 therefore depend only on the equilibrium labels of *A*, as required.

It follows from Corollary 1 that coarse-graining can be carried out on an equilibrium graph, *A*, by choosing any graph *G* for which 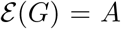 and carrying out the coarse-graining procedure described above on *G*. This justifies the coarse-graining construction described in the main text.

### COARSE GRAINING THE ALLOSTERY GRAPH

#### Proof of Eq.3

As described in the main text and Fig.3, the coarse-grained allostery graph, 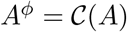, is defined using the partition of *A* by its horizontal subgraphs, *A_s_*, where *S* runs through all binding subsets, *S* ⊆ {1, ⋯, *n*}. *A^ϕ^* has the same structure of vertices and edges as any of the binding subgraphs, *A^c_k_^*, and is indexed in the same way by the binding subsets, *S*. Scheme 1 shows an example, which illustrates the calculations undertaken in this section.

Consider the reversible edge in *A^ϕ^*, *S* ⇌ *S* ∪ {*i*}, where *i* ∉ *S*. This reversible edge effectively arises from the binding and unbinding of ligand at site *i*. According to Eq.13, its effective association constant, 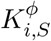 should satisfy,

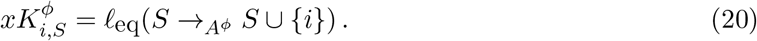

Since *A* is at thermodynamic equilibrium, we can make use of the formula in Corollary 1 to rewrite this as,

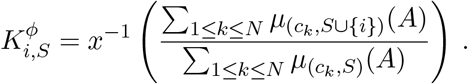

Eqs.8 and 13 tell us that 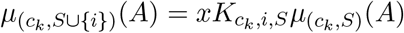, so that, after rearranging,

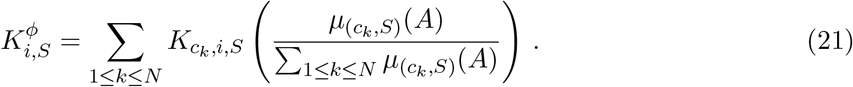

We can now appeal to Eqs.12 and 14 to rewrite the term in brackets on the right as,

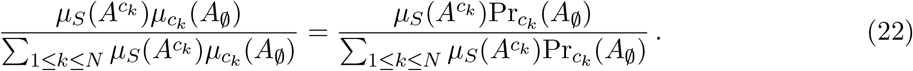

At this point, it will be helpful to introduce the following notation. If *G* is any equilibrium graph and *u* : *v*(*G*) → **R** is any real-valued function defined on the vertices of *G*, let 〈*u*〉 denote the average of *u* over the steady-state probability distribution of *G*,

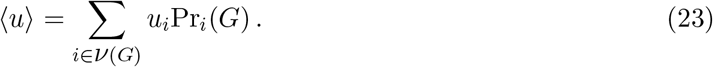

With this notation in hand, we can rewrite the denominator in Eq.22 as 〈*μ_S_*(*A^c_k_^*)), where, from now on, averages will be taken over the steady-state probability distribution of the horizontal subgraph of empty conformations, 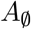 (Scheme 1, bottom). Inserting this expression back into Eq.21 and rearranging, we obtain a formula for the effective association constant as a ratio of averages,

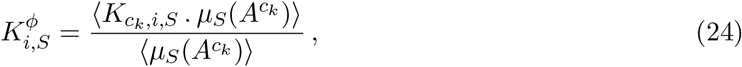

Here, the “dot” signifies a product to make the formula easier to read. Scheme 1 demonstrates this calculation. Recall from the main text that higher-order cooperativities (HOCs) are defined by normalising to the empty binding subset, so that 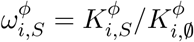. Furthermore, since the reference vertex of the vertical subgraphs, *A^c_k_^*, is taken to be the empty binding subset, 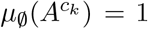. It follows that the effective HOCs are given by,

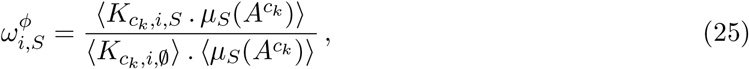

which gives Eq.3.

#### Elementary properties of effective HOCs

The main text describes three elementary properties of effective HOCs which follow from Eq.25. The only quantity in Eq.25 which involves the ligand concentration, *x*, is *μ_s_*(*A^c_k_^*). It follows from Eq.8 that this quantity is a monomial in *x* of the form *ax^p^*, where *a* does not involve *x* and *p* = #(*S*). In particular, *x^p^* does not depend on the conformation *c_k_*. It follows that *x^p^* can be extracted from the averages in Eq.25 and cancelled between the numerator and denominator. Hence, 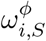 is independent of *x*. If 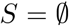, then *μ_s_*(*A^c_k_^*) = 1 for all 1 ≤ *k* ≤ *N* and it follows from Eq.25 that 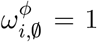. Finally, if there is only one conformation c_1_, the averages in Eq.25 collapse and *μ_S_*(*A*^*c*_1_^) cancels above and below, so that 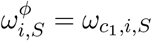, as required.

#### Generalised Monod-Wyman-Changeux formula

The original MWC formula calculates the binding curve, or fractional saturation, of the two-conformation model as a function of ligand concentration x (Monod et al., 1965). Here, we do the same for an arbitrary allostery graph, *A*. Let *s* = #(*S*). The fractional saturation of *A* is given by the average binding,

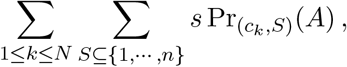

normalised to the number of binding sites, *n*. By the coarse graining formula in Eq.18, we can rewrite the fractional saturation as,

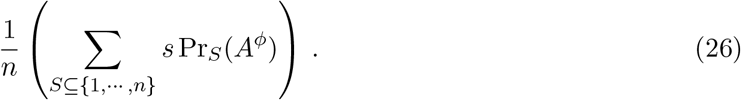

The probability, Pr*_S_*(*A^ϕ^*), can be calculated using Eq.10, which requires the quantities *μ_S_*(*A^ϕ^*). These can in turn be calculated by the path formula in Eq.8. We can choose the path in *A^ϕ^* to use the independent parameters introduced above. Let *S* = {*i*_1_, ⋯, *i_s_*}, where *i*_1_ < ⋯ < *i_s_*. Making use of Eq.20, we see that

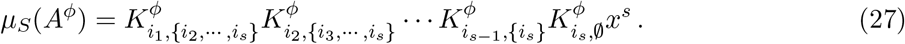

Eq.27 can be rewritten in terms of the non-dimensional effective HOCs but it is simpler for our purposes to use instead the effective association constants, 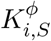. The dependence on x in Eq.27 shows that average binding is given by the logarithmic derivative of the partition function, Ψ(*A^ϕ^*), so the fractional saturation can be written,

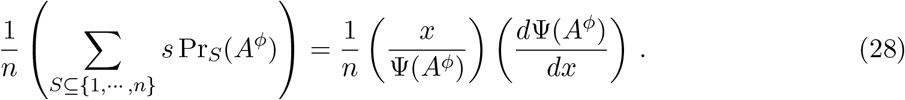

With this in mind, Eq.27 shows that the partition function can be written as a polynomial in *x*,

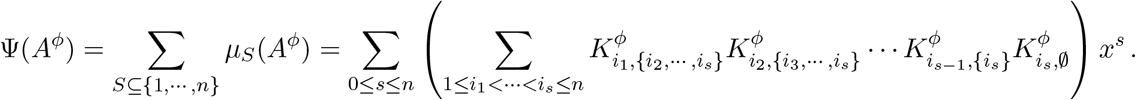

Finally, the 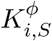 can be determined as averages over the horizontal subgraph of empty conformations using Eq.24. In this way, the fractional saturation in Eq.28 is ultimately determined by the independent parameters of *A*, giving rise thereby to a generalised Monod-Wyman-Changeux formula that is valid for any allostery graph. We explain below how the classical MWC formula is recovered using this procedure.

### EFFECTIVE HOCs FOR MWC-LIKE MODELS

#### Proof of Eq.4 and related work

Let *A* be an allostery graph with ligand binding to *n* sites which are independent and identical in each conformation. Because of independence, *ω_c_k_,i,S_* = 1, so that 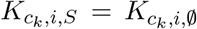 does not depend on *S*; because the sites are identical, *K_c_i_,i,S_* does not depend on *i*. Hence, we may write, *K_c_k_,i,S_* = *K_c_k__* and the labels on the binding edges of the vertical subgraph *A^c_k_^* are all given by *K_c_k__*. It follows from Eq.8 that *μ_S_*(*A^c_k_^*) = (*K_c_k__*)*^s^*, where *s* = #(*S*). Eq.25 then tells us that 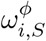 also depends only on *s*, so that we can write it as 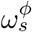, and Eq.25 simplifies to,

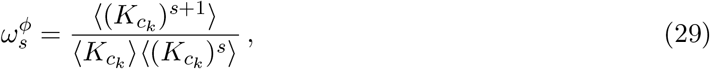

which gives Eq.4.

If we consider the effective association constant instead of the effective HOC, then, with the same assumptions as above, Eq.24 tells us that,

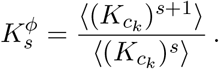

Suppose that only two conformations, *R* and *T*, are present. Let ℓ_eq_(*c_R_* → *c_T_*) = *L* and write *K_c_T__* and *K_c_R__* as *K_T_* and *K_R_*, respectively. Then, for any random variable on conformations, *X_c_k__*, the average is given by, 〈*X_c_k__*〉 = (*X_c_R__* + *X_c_T__L*)/(1 + *L*). Hence,

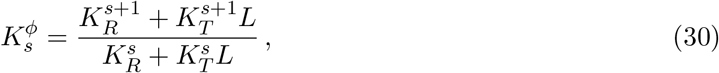

which is the formula for the (*s* + 1)-th “intrinsic binding constant” given by Gruber and Horovitz in (Gruber and Horovitz, 2018, Eq.(2.10)). In their analysis, the word “intrinsic” means what we would call “effective”.

We can use Eq.30 to work out what the generalised MWC formula derived above yields for the classical MWC model. Substituting Eq.30 in Eq.27, the intermediate terms in the product cancel out to leave,

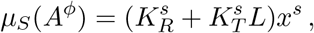

in which the right-hand side depends only on *s* = #(*S*). Collecting together subsets of the same size, the partition function of *A^ϕ^* may be written as,

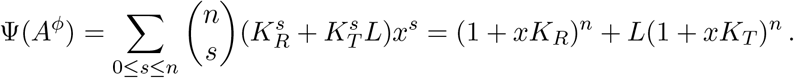

It then follows from Eq.28 that the fractional saturation is given by,

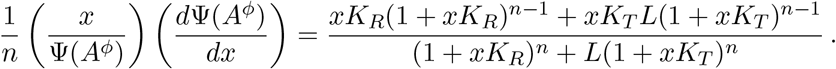

If we set *α* = *xK_R_* and *cα* = *xK_T_* this gives, for the fractional saturation,

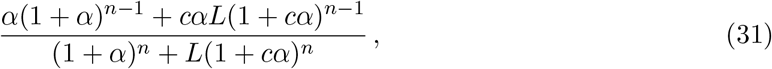

which recovers the classical Monod-Wyman-Changeux formula in the notation of (Monod et al., 1965, Eq.2).

#### Proof of Eq.5

The following result is unlikely not to be known in other contexts but we have not been able to find mention of it.

##### Lemma 3

*Suppose that X is a positive random variable, X* > 0, *over a finite probability distribution. If s* ≥ 1, *the following moment inequality holds*,

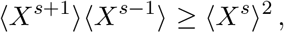

*with equality if, and only if, X is constant over the distribution*.

**Proof:** Suppose that the states of the probability space are indexed by 1 ≤ *i* ≤ *m* and that *p_i_* denotes the probability of state *i*. Then,

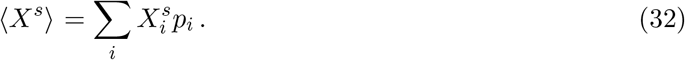

The quantity *α_s_* = 〈*X*^*s*+1^〉〈*X*^*s*−1^〉 – 〈*X^s^*〉^2^ can then be written as,

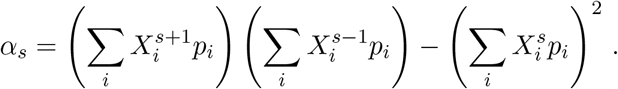

Collecting together terms in *p_i_p_j_*, we can rewrite this as

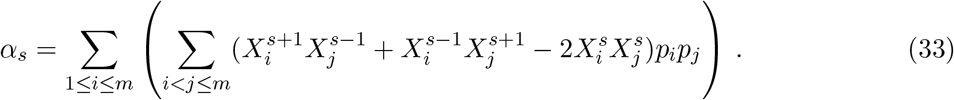

Note that the terms corresponding to *i* = *j* yield 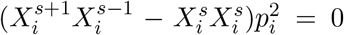 and so do not contribute to Eq.33. Choose any pair 1 ≤ *i* ≤ *m* and *i* < *j* ≤ *m* and let *X_j_* = *μX_i_*. Then, the coefficient of *p_i_p_j_* in Eq.33 becomes,

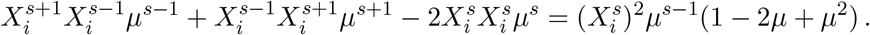

Now, 1 – 2*μ* + *μ*^2^ = (*μ* – 1)^2^ ≥ 0 for *μ* ∈ **R**, with equality if, and only if, *μ* = 1. Since *X* > 0 by hypothesis, *μ* > 0, so the coefficient of *p_i_p_j_* is positive unless *μ* = 1. Hence, *α_s_* > 0 unless *X_i_* = *X_j_* whenever 1 ≤ *i* ≤ *m* and *i* < *j* ≤ *m*, which means that *X* is constant over the distribution. Of course, if *X* is constant, then clearly *α_s_* = 0 for all *s* ≥ 1. The result follows.

##### Corollary 2

*If A is an MWC-like allostery graph, its effective HOCs satisfy*,

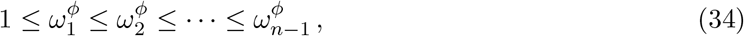

*with equality at any stage if, and only if, K_c_k__ is constant over* 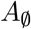.

**Proof:** It follows from Eq.29 that we can rewrite the effective HOCs recursively as,

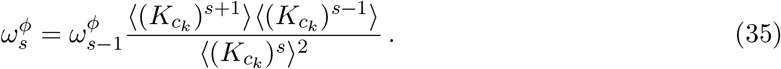

Since 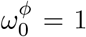, the result follows by recursively applying Lemma 3 to *X* = *K_c_k__* > 0. Eq.34 gives Eq.5.

#### Negative effective cooperativity

We consider an allostery graph *A* with two conformations and two sites, in which binding is independent but not identical, so that the association constants differ between sites. Let 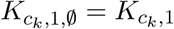 and 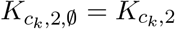, for *k* = 1,2. Since the sites are independent, *ω*_*c_k_*, 1, {2}_ = 1, so that *K*_*c_k_*, 1, {2}_ = *K*_*c_k_*, 1_, for *k* = 1,2. It follows from Eq.8—see also Scheme 1—that

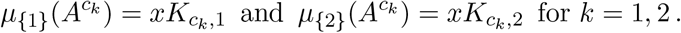

Let λ be the single equilibrium label in the horizontal subgraph of empty conformations,

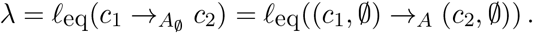

It follows from Eqs.8 and 10—see also the similar calculation in Scheme 1—that 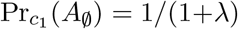 and 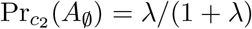. We know from Eq.25 that, and using the identifications above, we see that,

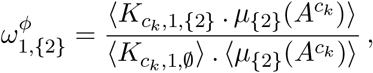

and using the identifications above, we see that,

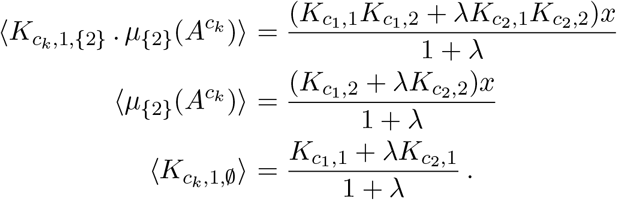

Substituting and simplifying, we find that,

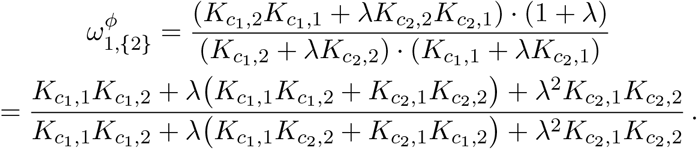

The first and last terms are the same in the numerator and denominator, so it follows that 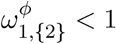 if, and only if,

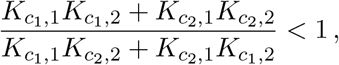

which is to say,

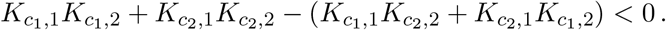

The left-hand side factors to give,

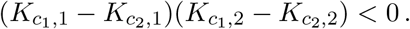

We see that negative cooperativity arises if, and only if, the sites have opposite patterns of association constants in the two conformations.

### FLEXIBILITY OF ALLOSTERY

#### The integrative flexibility theorem

Some preliminary notation is needed. Recall that if *X* is a finite set—typically, a subset of {1, ⋯, *n*}—then #(*X*) will denote the number of elements in *X*. If *X* and *Y* are sets, then *X*\*Y* will denote the complement of *Y* in *X*, *X*\*Y* = {*i* ∈ *X*, *i* ∉ *Y*}. Recall also the “little o” notation: 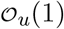 will stand for any quantity which depends on *u* and for which 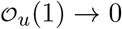 as *u* → 0. For instance, *Au* + *Bu*^2^ is 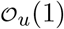 but (*Au* + *Bu*^2^)/*u* is 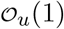 if, and only if, *A* = 0. This notation allows concise expression of complicated expressions which vanish in the limit as *u* → 0. Note that *f* (*u*) → *A* as *u* → 0 if, and only if, 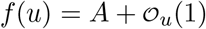, which is a useful trick for simplifying *f*.

##### Theorem 1

*Suppose n* ≥ 1 *and choose* 2*^n^* – 1 *arbitrary positive numbers*

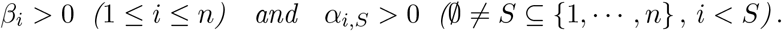

*Given any ε* > 0 *and δ* > 0, *there exists an allosteric conformational ensemble, which has no intrinsic HOC in any conformation, such that*

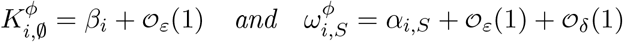

*for all corresponding values of i and S*.

**Proof:** We will construct an allostery graph *A* whose conformations are indexed by subsets *T* ⊆ {1, ⋯, *n*} and denoted *c_T_* (Scheme 2). Both conformations and binding subsets will then be indexed by subsets of {1, ⋯, *n*}. They should not be confused with each other. The reference vertex of *A* is 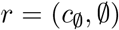. For the horizontal subgraph of empty conformations, 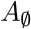, let 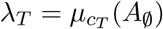. It follows from Eq.8, using *μ* in place of *ρ*, that the *λ_c_T__* determine the equilibrium labels of 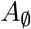. Keeping in mind that 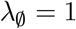, the λ*_T_* form a set of 2*^n^* – 1 independent parameters for 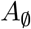, as explained above. The partition function of 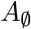 is then given by 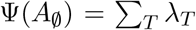 and the steady-state probabilities are given by 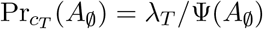 (Eq.12).

**Scheme 2:**
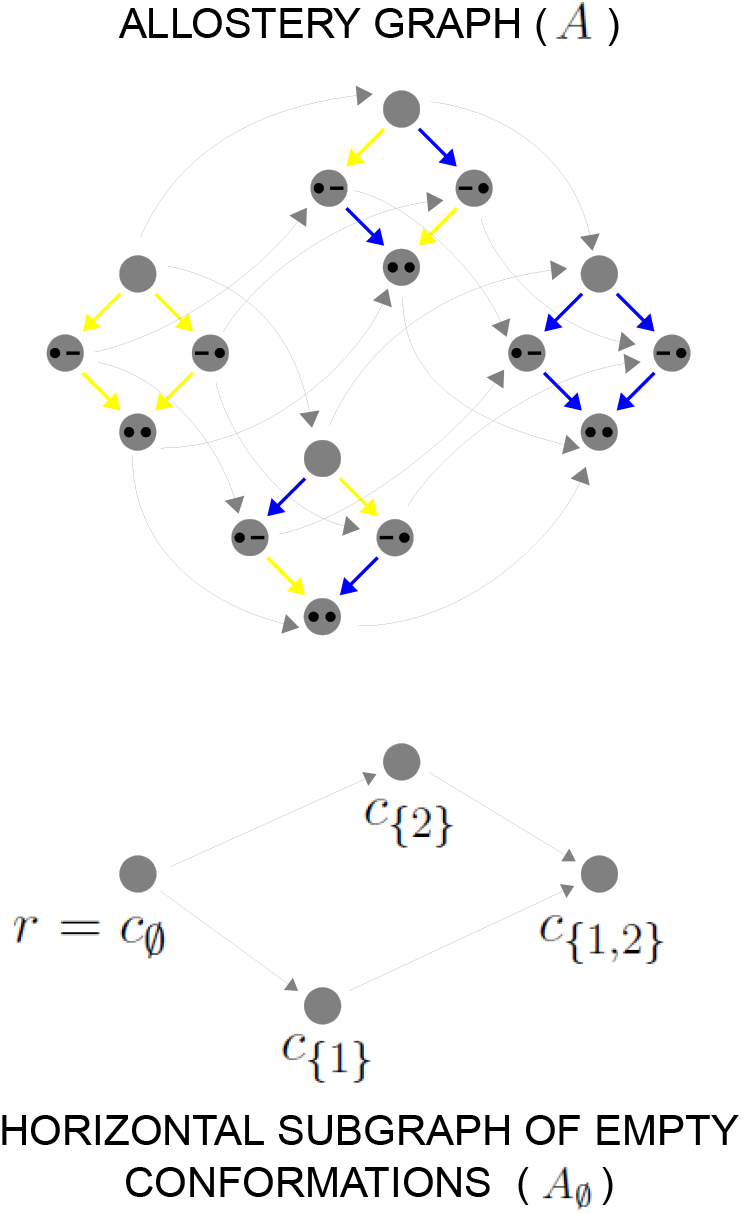
Example allostery graph for Theorem 1. There are *n* = 2 sites and 2^2^ = 4 conformations, giving a 16-vertex allostery graph (top), with similar conventions and format as in Scheme 1. The yellow vertical binding edges carry a factor of *ε* in their equilibrium labels, while the blue vertical binding edges do not, as specified by Eq.36. The horizontal subgraph of empty conformations is shown at the bottom, with conformations indexed by subsets of {1, 2}.

The other parameters will be *n* quantities from which we will construct the association constants in each conformation. Let *κ*_1_, ⋯, *κ_n_* > 0 be positive quantities whose values we will subsequently choose. We assume that all intrinsic HOCs are 1 and, for any binding microstate *S* ⊆ {1, ⋯, *n*}, we set,

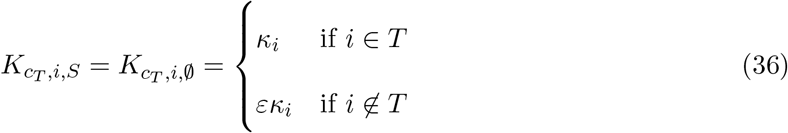

Scheme 2 illustrates Eq.36 for the case *n* = 2. If *c_T_* is a conformation and *S* ⊆ {1, ⋯, *n*} is a binding microstate, it follows from Eq.36 that,

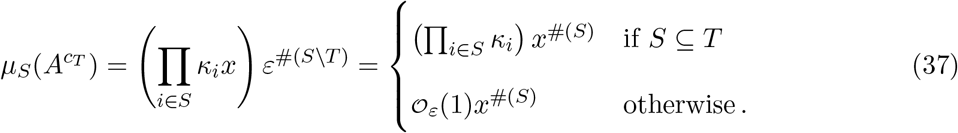

After coarse-graining, we can calculate effective association constants and effective HOCs using the formulas in Eqs.24 and 25. It follows from Eq.24 and Eq.36 that the effective bare association constants are,

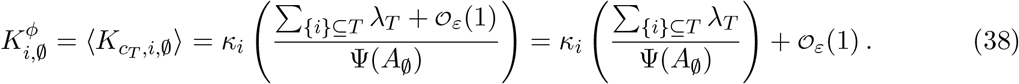

For the last step, we have used the trick mentioned above of letting *ε* → 0 to determine the term independent of *ε*. We have written {*i*} ⊆ *T* in place of the equivalent *i* ∈ *T* for consistency with formulas which will appear below. Note that the association constants in Eq.38 are linear in the *κ_i_*, whose values are therefore readily determined once the λ*_T_* have been chosen. Now let *S* be a binding microstate and *i* ∉ *S*. Using Eq.24 and Eqs.36 and 37,

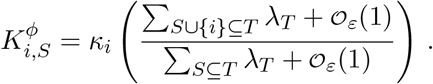

Letting *ε* → 0, we can use the trick above to rewrite this as,

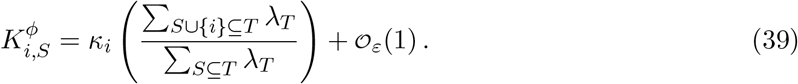

Note that Eq.39 simplifies to Eq.38 when 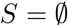. It follows from Eq.38 and Eq.39, using the same trick to reorganise the terms which are 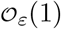, that the effective HOCs are,

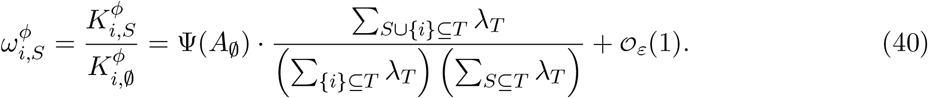

We see that the effective HOCs are independent of the quantities *κ_i_* and depend only on the parameters, λ*_T_*, of the horizontal subgraph 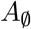.

We can now specify the λ*_T_*. Of course, 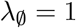. If *T* = {*i*_1_, ⋯, *i_k_*}, where *i*_1_ < *i*_2_ < ⋯ < *i_k_*, we set,

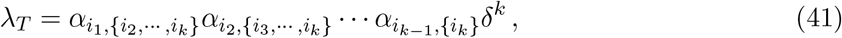

where each of the *α* quantities is given by hypothesis. In particular, if *T* = {*j*}, then λ_T_ = *δ*. Note that the exponent of *δ* depends only on the size of *T* and not on which elements *T* contains. For the example in Scheme 2, λ_{1}_ = λ_{2}_ = *δ* and λ_{1,2}_ = *α*_1,{2}_*δ*^2^.

It follows from Eq.41 that, given any *X* ⊆ {1, ⋯, *n*},

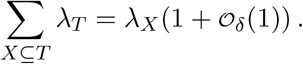

Using this, we see that the main term in Eq.40 has the form,

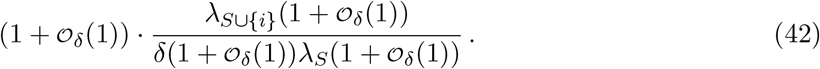

It follows from Eq.41 that, when 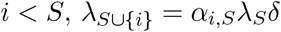, so using the trick above for reorganising the 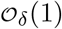 terms, we can rewrite Eq.42 as 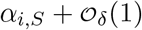. Substituting back into Eq.40, we see that, when *i* < *S*,

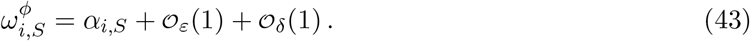

With the choice of λ*_T_* given by Eq.41, we can return to Eq.38 and define,

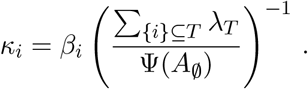

Substituting back into Eq.38, we see that,

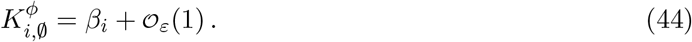

The result follows from Eqs.43 and 44.

In respect of the dimensional argument made in the main text, the allostery graph constructed in Theorem 1 has 2*^n^* – 1 free parameters for 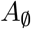 and the *κ*_1_, ⋯, *κ_n_* are a further *n* free parameters, giving 2*^n^* – 1 + *n* free parameters in total. This is more than the minimal required number of 2*^n^* – 1 but not by much. It remains an interesting open question whether a conformational ensemble can be constructed, perhaps with more free parameters, which gives the effective HOCs exactly, rather than only approximately.

#### Construction of Fig.4

We implemented in a Mathematica notebook the proof strategy in Theorem 1 for any number *n* of sites. The notebook takes as input parameters the *β_i_* and the *α_i,S_* for *i* < *S* in the statement of the theorem, along with specified values for the quantities *ε* and *δ*. It produces as output the effective bare association constants, 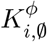, and effective HOCs, 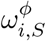 for *i* < *S*, as given by Theorem 1. The values of *ϵ* and *δ* can then be adjusted so that the calculated 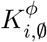 and 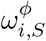 are as close as required to the *β_i_* and *α_i,S_*. The notebook is available on request.

Fig.4 shows the results from using this notebook on three examples, chosen by hand to illustrate different patterns of effective bare association constants and effective HOCs. The actual numerical values are listed below. The colour names used here refer to the colour code for the three examples in Fig.4. The maximum error was calculated as the larger of 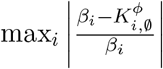 and 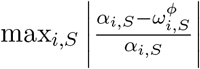. The quantities *δ* and *ε* were adjusted to make the maximum error less than 0.01.

In Fig.4, the conformations are indexed *c*_1_, ⋯, *c*_16_. These indices enumerate the conformations *c_T_*, for subsets *T* ⊆ {1, ⋯, 4}, used in the proof of Theorem 1, according to the following ordering: the subsets are arranged in blocks of increasing size starting from 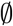 and within each block they are arranged in increasing lexicographic order when the elements in each set are listed in increasing numerical order. Accordingly, the relationship between numerical indices and subsets is as specified below.

| | maroon<br>$\delta = 10^{-7}, \varepsilon = 10^{-12}$ | | orange<br>$\delta = 10^{-7}, \varepsilon = 10^{-14}$ | | red<br>$\delta = 10^{-7}, \varepsilon = 10^{-16}$ | |
| --- | --- | --- | --- | --- | --- | --- |
| $i$ | $\beta_i$ | $K_{i,\emptyset}^\phi$ | $\beta_i$ | $K_{i,\emptyset}^\phi$ | $\beta_i$ | $K_{i,\emptyset}^\phi$ |
| 1 | 1.5777 | 1.5776 | 0.031353 | 0.031353 | 0.21257 | 0.21257 |
| 2 | 24.013 | 24.014 | 0.011104 | 0.011104 | 0.84301 | 0.84301 |
| 3 | 89.958 | 89.959 | 13.195 | 13.195 | 9.8514 | 9.8514 |
| 4 | 0.015685 | 0.015685 | 52.437 | 52.437 | 27.000 | 27.000 |
| $i, S$ | $\alpha_{i,S}$ | $\omega_{i,S}^\phi$ | $\alpha_{i,S}$ | $\omega_{i,S}^\phi$ | $\alpha_{i,S}$ | $\omega_{i,S}^\phi$ |
| 1, {2} | 0.084815 | 0.0848456 | 1.0801 | 1.0801 | 50.455 | 50.454 |
| 1, {3} | 3.7432 | 3.7432 | 34.768 | 34.768 | 0.016359 | 0.016401 |
| 1, {4} | 0.044245 | 0.044264 | 0.032668 | 0.032669 | 0.60018 | 0.60018 |
| 2, {3} | 30.240 | 30.239 | 4.0683 | 4.0683 | 7.2944 | 7.2944 |
| 2, {4} | 0.074064 | 0.074083 | 1.5098 | 1.5098 | 0.010809 | 0.010809 |
| 3, {4} | 9.2687 | 9.2685 | 0.025183 | 0.025184 | 0.012613 | 0.012613 |
| 1, {2, 3} | 4.0933 | 4.0933 | 0.31238 | 0.31238 | 57.783 | 57.783 |
| 1, {2, 4} | 15.687 | 15.683 | 0.70016 | 0.70016 | 0.025618 | 0.025623 |
| 1, {3, 4} | 0.013335 | 0.013349 | 0.13042 | 0.13056 | 4.4450 | 4.4450 |
| 2, {3, 4} | 0.082851 | 0.082892 | 2.5235 | 2.5235 | 0.13584 | 0.13584 |
| 1, {2, 3, 4} | 6.5843 | 6.5825 | 0.017404 | 0.017407 | 0.063587 | 0.063833 |
| max. error | 0.00105 |  | 0.00105 |  | 0.00386 |  |

|  |  |  |  |  |  |  |  |  |  |  |
| --- | --- | --- | --- | --- | --- | --- | --- | --- | --- | --- |
| $c_1$ | $c_2$ | $c_3$ | $c_4$ | $c_5$ | $c_6$ | $c_7$ | $c_8$ | $c_9$ | $c_{10}$ | $c_{11}$ |
| $c_\emptyset$ | $c_{\{1\}}$ | $c_{\{2\}}$ | $c_{\{3\}}$ | $c_{\{4\}}$ | $c_{\{1,2\}}$ | $c_{\{1,3\}}$ | $c_{\{1,4\}}$ | $c_{\{2,3\}}$ | $c_{\{2,4\}}$ | $c_{\{3,4\}}$ |
| | | $c_{12}$ | $c_{13}$ | $c_{14}$ | $c_{15}$ | $c_{16}$ | | | | |
| | | $c_{\{1,2,3\}}$ | $c_{\{1,2,4\}}$ | $c_{\{1,3,4\}}$ | $c_{\{2,3,4\}}$ | $c_{\{1,2,3,4\}}$ | | | | |

The binding curves for each example (Fig.4, right) show the dependence on concentration of average binding to site *i* (coloured curves), which can be written in terms of the coarse-grained graph, 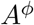, in the form,

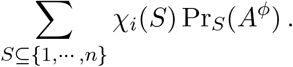

Here, *χ_i_*(*S*) is the indicator function for *i* being in *S*,

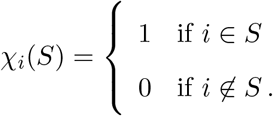

Since the size of *S*, which was denoted by *s* above, is given by 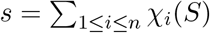, we see from Eq.26 that the fractional saturation (Fig.4, right, black curves) is the sum of the average bindings over all sites, normalised to the number of sites, *n*.

It should be noted that the particular method used to prove Theorem 1 necessarily yields an exponentially wide range of parameter values, as is evident in Fig.4. It is another interesting open question as to whether alternative constructions are possible, with more constrained parameter values.

## Author Contributions

All authors formulated the project and performed research. J.G. wrote the paper with the assistance of all authors.

## Acknowledgments

JWB and JG were supported by US National Science Foundation (NSF) Award #1462629. RMC was supported by US National Institutes of Health award #GM122928 and by EMBO Fellowship ALTF683-2019. FW was supported by the James S. McDonnell Foundation and NSF Graduate Research Fellowship #DGE1144152.

## References

Ahsendorf T, Wong F, Eils R, Gunawardena J. A framework for modelling gene regulation which accommodates non-equilibrium mechanisms. BMC Biol. 2014; 12:102.

Allen BL, Taatjes DJ. The Mediator complex: a central integrator of transcription. Nat Rev Mol Cell Biol. 2015; 16:155–66.

Bacic L, Sabantsev A, Deindl S. Recent advances in single-molecule fluorescence microscopy render structural biology dynamic. Curr Opin Struct Biol. 2020; 65:1–8.

Benabdallah NS, Bickmore WA. Regulatory domains and their mechanisms. Cold Spring Harb Symp Quant Biol. 2015; 80:45–51.

Berlow RB, Dyson HJ, Wright PE. Expanding the paradigm: intrinsically disordered proteins and allosteric regulation. J Mol Biol. 2018; 430:2309–20.

Biddle JW, Nguyen M, Gunawardena J. Negative reciprocity, not ordered assembly, underlies the interaction of Sox2 and Oct4 on DNA. eLife. 2019; 8:e410172018.

Bolt CC, Duboule D. The regulatory landscapes of developmental genes. Development. 2020; 147:dev171736.

Changeux JP. The feedback control mechanism of biosynthetic L-threonine deaminase by L-isoleucine. Cold Spring Harb Symp Quant Biol. 1961; 26:389–401.

Changeux JP. 50 years of allosteric interactions: the twists and turns of the models. Nat Rev Mol Cell Biol. 2013; 14:1–11.

Changeux JP, Christopoulos A. Allosteric modulation as a unifying mechanism for receptor function and regulation. Cell. 2016; 166:1084–102.

Chong S, Dugast-Darzacq C, Liu Z, Dong P, Dailey GM, Cattoglio C, Heckert A, Banala S, Lavis L, Darzacq X, Tjian R. Imaging dynamic and selective low-complexity domain interactions that control gene transcription. Science. 2018; 361:eaar2555.

Clark S, Myers JB, King A, Fiala R, Novacek J, Pearce G, Heierhorst J, Reichow SL, Barbar EJ. Multivalency regulates activity in an intrinsically disordered transcription factor. eLife. 2018; 7:e36258.

Cooper A, Dryden DT. Allostery without conformational change. A plausible model. Eur Biophys J. 1984; 11:103–9.

Dasgupta T, Croll DH, Owen JA, Vander Heiden MG, Locasale JW, Alon U, Cantley LC, Gunawardena J. A fundamental trade off in covalent switching and its circumvention by enzyme bifunctionality in glucose homeostasis. J Biol Chem. 2014; 289:13010–25.

Demir Ö, Leong PU, Amaro RE. Full-length p53 tetramer bound to DNA and its quaternary dynamics. Oncogene. 2017; 36:1451–60.

Dodd IB, Shearwin KE, Perkins AJ, Burr T, Hochschild A, Egan JB. Cooperativity in long-range gene regulation by the λ CI repressor. Genes and Development. 2004; 18:344–54.

Dyson HJ, Wright PE. Role of intrinsic protein disorder in the function and interactions of the transcriptional coactivators CREB-binding protein (CBP) and p300. J Biol Chem. 2016; 291:6714–22.

Edelman LB, Fraser P. Transcription factories: genetic programming in three dimensions. Curr Opin Genet Dev. 2012; 22:110–14.

Ehlert FJ. Cooperativity has empirical and ultimate levels of explanation. Trends Pharmacol Sci. 2016; 37:620–3.

Estrada J, Wong F, DePace A, Gunawardena J. Information integration and energy expenditure in gene regulation. Cell. 2016; 166:234–44.

Frauenfelder H, Sligar SG, Wolynes PG. The energy landscapes and motions of proteins. Science. 1991; 254:1598–603.

Freidlin MI, Wentzell AD. Random perturbations of dynamical systems. 3 ed. Heidleberg, Germany: Springer; 2012.

Furlong EEM, Levine M. Developmental enhancers and chromosome topology. Science. 2018; 361:1341–5.

Ganser LR, Kelly ML, Herschlag D, Al-Hashimi HM. The roles of structural dynamics in the cellular functions of RNAs. Nat Rev Mol Cell Biol. 2019; 20:474–89.

Gerhart J. From feedback inhibition to allostery: the enduring example of aspartate transcar-bamoylase. FEBS J. 2014; 281:612–20.

Gruber R, Horovitz A. Unpicking allosteric mechanisms of homo-oligomeric proteins by determining their successive ligand binding constants. Phil Trans R Soc B. 2018; 373:20170176.

Gunawardena J. A linear framework for time-scale separation in nonlinear biochemical systems. PLoS ONE. 2012; 7:e36321.

Gunawardena J. Time-scale separation: Michaelis and Menten’s old idea, still bearing fruit. FEBS J. 2014; 281:473–88.

Henzler-Wildman K, Kern D. Dynamic personalities of proteins. Nature. 2007; 450:964–72.

Hilser VJ, Wrabl JO, Motlagh HN. Structural and energetic basis of allostery. Annu Rev Biophys. 2012; 41:585–609.

Kalo A, Kanter I, Shraga A, Sheinberger J, Tzemach H, Kinor N, Singer RH, Lionnet T, Shav-Tal Y. Cellular levels of signaling factors are sensed by β-actin alleles to modulate transcriptional pulse intensity. Cell Rep. 2015; 11:419–32.

Kim S, Brostrmer E, Xing D, Jin J, Chong S, Ge H, Wang S, Gu C, Yang L, Gao YQ, Su XD, Sun Y, Xie XS. Probing allostery through DNA. Science. 2013; 339:816–9.

Knoverek CR, Amarasinghe GK, Bowman GR. Advanced methods for accessing protein shape-shifting present new therapeutic opportunities. Trends Biochem Sci. 2019; 44:351–64.

Kornev AP, Taylor SS. Dynamics-driven allostery in protein kinases. Trends Biochem Sci. 2015; 40:628–47.

Koshland DE, Hamadani K. Proteomics and models for enzyme cooperativity. J Biol Chem. 2002; 277:46841–4.

Koshland DE, Nemethy G, Filmer D. Comparison of experimental binding data and theoretical models in proteins containing subunits. Biochemistry. 1966; 5:365–85.

LeVine MV, Weinstein H. AIM for allostery: using the Ising model to understand information processing and transmission in allosteric biomolecular systems. Entropy. 2015; 17:2895–918.

Lewis BA. Understanding large multiprotein complexes: applying a multiple allosteric networks model to explain the function of the Mediator transcription complex. J Cell Sci. 2010; 123:159–63.

Lewis JS, Costa A. Caught in the act: structural dynamics of replication origin activation and fork progression. Biochem Soc Trans. 2020; doi:10.1042/BST20190998.

Li J, Dong A, Saydaminova K, Chang H, Wang G, Ochiai H, Yamamoto T, Pertsinidis A. Single-molecule nanoscopy elucidates RNA polymerase II transcription at single genes in live cells. Cell. 2019; 178:491–506.

Lin Y, Sohn CH, Dalal CK, Cai L, Elowitz MB. Combinatorial gene regulation by modulation of relative pulse timing. Nature. 2015; 527:54–8.

Liu J, Perumal NB, Oldfield CJ, Su EW, Uversky VN, Dunker AK. Intrinsic disorder in transcription factors. Biochemistry. 2006; 45:6873–88.

Lorimer GH, Horovitz A, McLeish T. Allostery and molecular machines. Phil Trans R Soc B. 2018; 373:20170173.

Marco A, Meharena HS, Dileep V, Raju1 RM, Davila-Velderrain J, Zhang AL, Adaikkan C, Young JZ, Gao F, Kellis M, Tsai LH. Mapping the epigenomic and transcriptomic interplay during memory formation and recall in the hippocampal engram ensemble. Nat Neurosci. 2020; doi:10.1038/s41593-020-00717-0.

Martini JWR. A measure to quantify the degree of cooperativity in overall titration curves. J Theor Biol. 2017; 432:33–7.

Marzen S, Garcia HG, Phillips R. Statistical mechanics of Monod-Wyman-Changeux (MWC) models. J Mol Biol. 2013; 425:1433–60.

Mir M, Bickmore W, Furlong EEM, Narlikar G. Chromatin topology, condensates and gene regulation: shifting paradigms or just a phase? Development. 2019; 146:dev182766.

Mirny L. Nucleosome-mediated cooperativity between transcription factors. Proc Natl Acad Sci USA. 2010; 107:22534–9.

Mirzaev I, Bortz DM. Laplacian dynamics with synthesis and degradation. Bull Math Biol. 2015; 77:1013–45.

Mirzaev I, Gunawardena J. Laplacian dynamics on general graphs. Bull Math Biol. 2013; 75:2118–49.

Molina N, Suterb DM, Cannavoa R, Zollera B, Goticb I, Naef F. Stimulus-induced modulation of transcriptional bursting in a single mammalian gene. Proc Natl Acad Sci USA. 2013; 110:20563–8.

Monod J, Jacob F. General conclusions: teleonomic mechanisms in cellular metabolism, growth and differentiation. Cold Spring Harbor Symp Quant Biol. 1961; 26:389–401.

Monod J, Wyman J, Changeux JP. On the nature of allosteric transitions: a plausible model. J Mol Biol. 1965; 12:88–118.

Motlagh HN, Li J, Thompson EB, Hilser VJ. Interplay between allostery and intrinsic disorder in an ensemble. Biochem Soc Trans. 2012; 40:975–80.

Motlagh HN, Wrabl JO, Li J, Hilser VJ. The ensemble nature of allostery. Nature. 2014; 508:331–9.

Noé F, Fischer S. Transition networks for modeling the kinetics of conformational change in macromolecules. Curr Opin Struct Biol. 2008; 18:154–62.

Nogales E, Fang J, Louder RK. Structural dynamics and DNA interaction of human TFIID. Transcription. 2017; 8:55–60.

Nussinov R, Tsai CJ, Ma B. The underappreciated role of allostery in the cellular network. Annu Rev Biophys. 2013; 42:169–89.

Park J, Estrada J, Johnson G, Ricci-Tam C, Bragdon M, Shulgina Y, Cha A, Gunawardena J, DePace AH. Dissecting the sharp response of a canonical developmental enhancer reveals multiple sources of cooperativity. eLife. 2019; 8:e41266.

Park M, Patel N, Keung AJ, Khalil AS. Engineering epigenetic regulation using synthetic read-write modules. Cell. 2019; 176:227–38.

Pauling L. The oxygen equilibrium of hemoglobin and its structural interpretation. Proc Natl Acad Sci USA. 1935; 21:186–91.

Peeters E, van Oeffelen L, Nidal M, Forterre P, Charlier D. A thermodynamic model of the cooperative interaction between the archaeal transcription factor Ss-LrpB and its tripartite operator DNA. Gene. 2013; 524:330–40.

Perutz MF. Stereochemistry of cooperative effects in haemoglobin. Nature. 1970; 228:726–34.

Portz B, Lu F, Gibbs EB, Mayfield JE, Mehaffey MR, Zhang YJ, Brodbelt JS, Showalter SA, Gilmour DS. Structural heterogeneity in the intrinsically disordered RNA polymerase II C-terminal domain. Nat Comm. 2017; 8:15231.

Robert CH, Decker H, Richey B, Gill SJ, Wyman J. Nesting: Hierarchies of allosteric interactions. Proc Natl Acad Sci USA. 1987; 84:1891–5.

Sabari BR, Dall’Agnese A, Boija A, Klein IA, Coffey EL, Shrinivas K, Abraham BJ, Hannett NM, Zamudio AV, Manteiga JC, Li CH, Guo YE, Day DS, Schuijers J, Vasile E, Malik S, Hnisz D, Lee TI, Cisse II, Roeder RG, et al. Coactivator condensation at super-enhancers links phase separation and gene control. Science. 2018; 361:eaar3958.

Schueler-Furman O, Wodak SJ. Computational approaches to investigating allostery. Curr Opin Struct Biol. 2016; 41:159–71.

Sengupta U, Strodel B. Markov models for the elucidation of allosteric regulation. Phil Trans Roy Soc B. 2018; 373:20170178.

Shi H, Rangadurai A, Assi HA, Roy R, Case DA, Herschlag D, Yesselman JD, Al-Hashimi HM. Rapid and accurate determination of atomistic RNA dynamic ensemble models using NMR and structure prediction. Nat Comm. 2020; 11:5531.

Smale ST, S E Plevy ASW, Zhou L, Ramirez-Carrozzi VR, Pope SD, Bhatt DM, Tong AJ. Toward an understanding of the gene-specific and global logic of inducible gene transcription. Cold Spring Harb Symp Quant Biol. 2013; 78:61–8.

Stroock DW. An Introduction to Markov Processes. 2nd ed. Graduate Texts in Mathematics, Berlin, Germany: Springer-Verlag; 2014.

Thal DM, Glukhova A, Sexton PM, Christopoulos A. Structural insights into G-protein coupled receptor allostery. Nature. 2018; 559:45–53.

Tsai CJ, Nussinov R. Gene-specific transcription activation via long-range allosteric shape-shifting. Biochem J. 2011; 439:15–25.

Tsai CJ, Nussinov R. A unified view of ’how allostery works’. PLoS Comput Biol. 2014; 10:e1003394.

Tzeng SR, Kalodimos CG. Protein dynamics and allostery: an NMR view. Curr Opin Struct Biol. 2011; 21:62–7.

Ullmann A. In memoriam: Jacques Monod (19101976). Genome Biol Evol. 2011; 3:1025–33.

Ventsel’ AD, Freidlin MI. On small random dynamical perturbations of dynamical systems. Russ Math Surv. 1970; 25:1–54.

Voss TC, Hager GL. Dynamic regulation of transcriptional states by chromatin and transcription factors. Nature Rev Genet. 2014; 15:69–81.

Wales DJ. Energy landscapes: calculating pathways and rates. Int Rev Phys Chem. 2006; 25:237–82.

Wodak SJ, Paci E, Dokholyan NV, Berezovsky IN, Horovitz A, Li J, Hilser VJ, Bahar I, Karan-icolas J, Stock G, Hamm P, Stote RH, Eberhardt J, Chebaro Y, Dejaegere A, Cecchini M, Changeux JP, Bolhuis PG, Vreede J, Faccioli P, et al. Allostery in its many disguises: from theory to applications. Structure. 2019; 27:566–78.

Wong F, Amir A, Gunawardena J. Energy-speed-accuracy relation in complex networks for biological discrimination. Phys Rev E. 2018; 98:012420.

Wong F, Dutta A, Chowdhury D, Gunawardena J. Structural conditions on complex networks for the Michaelis-Menten input-output response. Proc Natl Acad Sci USA. 2018; 115:9738–43.

Wong F, Gunawardena J. Gene regulation in and out of equilibrium. Annu Rev Biophys. 2020; 49:199–226.

Wrabl JO, Gu J, Liu T, Schrank TP, Whitten ST, Hilser VJ. The role of protein conformational fluctuations in allostery, function, and evolution. Biophys Chem. 2011; 159:129–41.

Wright PE, Dyson HJ. Intrinsically disordered proteins in cellular signalling and regulation. Nat Rev Mol Cell Biol. 2015; 16:18–29.

Yordanov P, Stelling J. Steady-state differential dose response in biological systems. Biophys J. 2018; 114:723–36.

Yordanov P, Stelling J. Efficient manipulation and generation of Kirchhoff polynomials for the analysis of non-equilibrium biochemical reaction networks. J Roy Soc Interface. 2020; 17:20190828.

